# Cellular deconstruction of inflamed synovium defines diverse inflammatory phenotypes in rheumatoid arthritis

**DOI:** 10.1101/2022.02.25.481990

**Authors:** Fan Zhang, Anna Helena Jonsson, Aparna Nathan, Kevin Wei, Nghia Millard, Qian Xiao, Maria Gutierrez-Arcelus, William Apruzzese, Gerald F. M. Watts, Dana Weisenfeld, Joyce B. Kang, Laurie Rumker, Joseph Mears, Kamil Slowikowski, Kathryn Weinand, Dana E. Orange, Javier Rangel-Moreno, Laura Geraldino-Pardilla, Kevin D. Deane, Darren Tabechian, Arnold Ceponis, Gary S. Firestein, Mark Maybury, Ilfita Sahbudin, Ami Ben-Artzi, Arthur M. Mandelin, Alessandra Nerviani, Felice Rivellese, Costantino Pitzalis, Laura B. Hughes, Diane Horowitz, Edward DiCarlo, Ellen M. Gravallese, Brendan F. Boyce, Accelerating Medicines Partnership Program: Rheumatoid Arthritis and Systemic Lupus Erythematosus (AMP RA/SLE) Network, Larry W. Moreland, Susan M. Goodman, Harris Perlman, V. Michael Holers, Katherine P. Liao, Andrew Filer, Vivian P. Bykerk, Deepak A. Rao, Laura T. Donlin, Jennifer H. Anolik, Michael B. Brenner, Soumya Raychaudhuri, Jennifer Albrecht, Jennifer L. Barnas, Joan M. Bathon, David L. Boyle, S. Louis Bridges, Debbie Campbell, Hayley L. Carr, Adam Chicoine, Andrew Cordle, Michelle Curtis, Patrick Dunn, Lindsy Forbess, Peter K. Gregersen, Joel M. Guthridge, Lionel B. Ivashkiv, Kazuyoshi Ishigaki, Judith A. James, Gregory Keras, Ilya Korsunsky, Amit Lakhanpal, James A. Lederer, Zhihan J. Li, Yuhong Li, Andrew McDavid, Nida Meednu, Ian Mantel, Mandy J. McGeachy, Karim Raza, Yakir Reshef, Christopher Ritchlin, William H. Robinson, Saori Sakaue, Jennifer A. Seifert, Melanie H. Smith, Dagmar Scheel-Toellner, Paul J. Utz, Michael H. Weisman, Zhu Zhu

**Author notes:** These authors contributed equally (Fan Zhang, Anna Helena Jonsson, Aparna Nathan, Kevin Wei, Nghia Millard). These authors jointly supervised this work (Deepak A. Rao, Laura T. Donlin, Jennifer H. Anolik, Michael B. Brenner, Soumya Raychaudhuri).

## Abstract

Rheumatoid arthritis (RA) is a prototypical autoimmune disease that causes destructive tissue inflammation in joints and elsewhere. Clinical challenges in RA include the empirical selection of drugs to treat patients, inadequate responders with incomplete disease remission, and lack of a cure. We profiled the full spectrum of cells in inflamed synovium from patients with RA with the goal of deconstructing the cell states and pathways characterizing pathogenic heterogeneity in RA. Our multicenter consortium effort used multi-modal CITE-seq, RNA-seq, and histology of synovial tissue from 79 donors to build a >314,000 single-cell RA synovial cell atlas with 77 cell states from T, B/plasma, natural killer, myeloid, stromal, and endothelial cells. We stratified tissue samples into six distinct cell type abundance phenotypes (CTAPs) individually enriched for specific cell states. These CTAPs demonstrate the striking diversity of RA synovial inflammation, ranging from marked enrichment of T and B cells (CTAP-TB) to a congregation of specific myeloid, fibroblast, and endothelial cells largely lacking lymphocytes (CTAP-EFM). Disease-relevant cytokines, histology, and serology metrics are associated with certain CTAPs. This comprehensive RA synovial atlas and molecular, tissue-based CTAP stratification reveal new insights into RA pathology and heterogeneity, which could lead to novel targeted-treatment approaches in RA.

## Introduction

Rheumatoid arthritis (RA) is a systemic autoimmune disease affecting up to 1% of the population^1^. It causes synovial joint tissue inflammation and extra-articular manifestations that lead to pain, damage, disability^2–5^. The clinical course of RA has been transformed by targeted therapeutics, including those aimed at TNF, IL-1, IL-6, B cells, T cell co-stimulation, and the JAK-STAT pathway^2,6^. Unfortunately, many patients are refractory to these therapies and do not achieve remission. Less than 25% of patients achieve an ACR70 response to any subsequent treatment after failing first-line therapies^7–9^. While current treatments can partially ameliorate disease activity, there is no cure. Thus, there is a clinical need for new RA treatment targets and an improved ability to predict patient-specific responses to treatment.

Genetic diversity and highly variable responses to targeted therapeutics suggest that RA may be a heterogeneous disease^10–13^. For example, patients who produce antibodies specific for cyclic citrullinated peptides (CCP) have different HLA and non-HLA susceptibility factors compared to CCP-negative patients^14^. However, genetic differences and clinical differences in disease duration or activity have not reliably predicted treatment response or druggable targets thus far^15–18^.

A more granular understanding of tissue inflammation and cell states may reveal synovial phenotypes that could inform prognosis and potentially identify new treatment targets. Encouragingly, preliminary clinical trials using histological or bulk RNA-seq analysis of tissue suggest treatment response may depend on tissue cellular composition^19,20^. We and others previously identified specific effector cell states in RA pathophysiology that represent promising treatment targets including pro-inflammatory *HBEGF^+^IL1B^+^* macrophages, *MERTK^+^* macrophages, *ITGAX^+^TBX21^+^* autoimmune-associated B cells (ABCs), *PDCD1^+^* peripheral helper T (T_PH_) cells, and *NOTCH3^+^* synovial fibroblasts^21–27^. We do not yet know whether this is a comprehensive list of disease-associated populations and if these disease-associated populations are present in every patient with RA.

To deconstruct the inflammatory cellular components of RA synovium, we analyzed cell-state composition in a diverse set of patients with clinically active RA. We sought to determine whether certain states are enriched only in certain subsets of patients. Since RA is a prototypical autoimmune disease that shares disease-associated tissue cell states^23,28–32^ and risk loci with other autoimmune diseases^33,34^, these analyses may offer insights into other diseases in which tissue inflammation is a hallmark.

## Results

To characterize RA patient heterogeneity, we utilized a multimodal single-cell synovial tissue pipeline to stratify tissue samples into distinct subgroups, characterize their associated cell states, and identify their clinical and histologic associations (**Figure 1A-D**).

**Figure 1.**
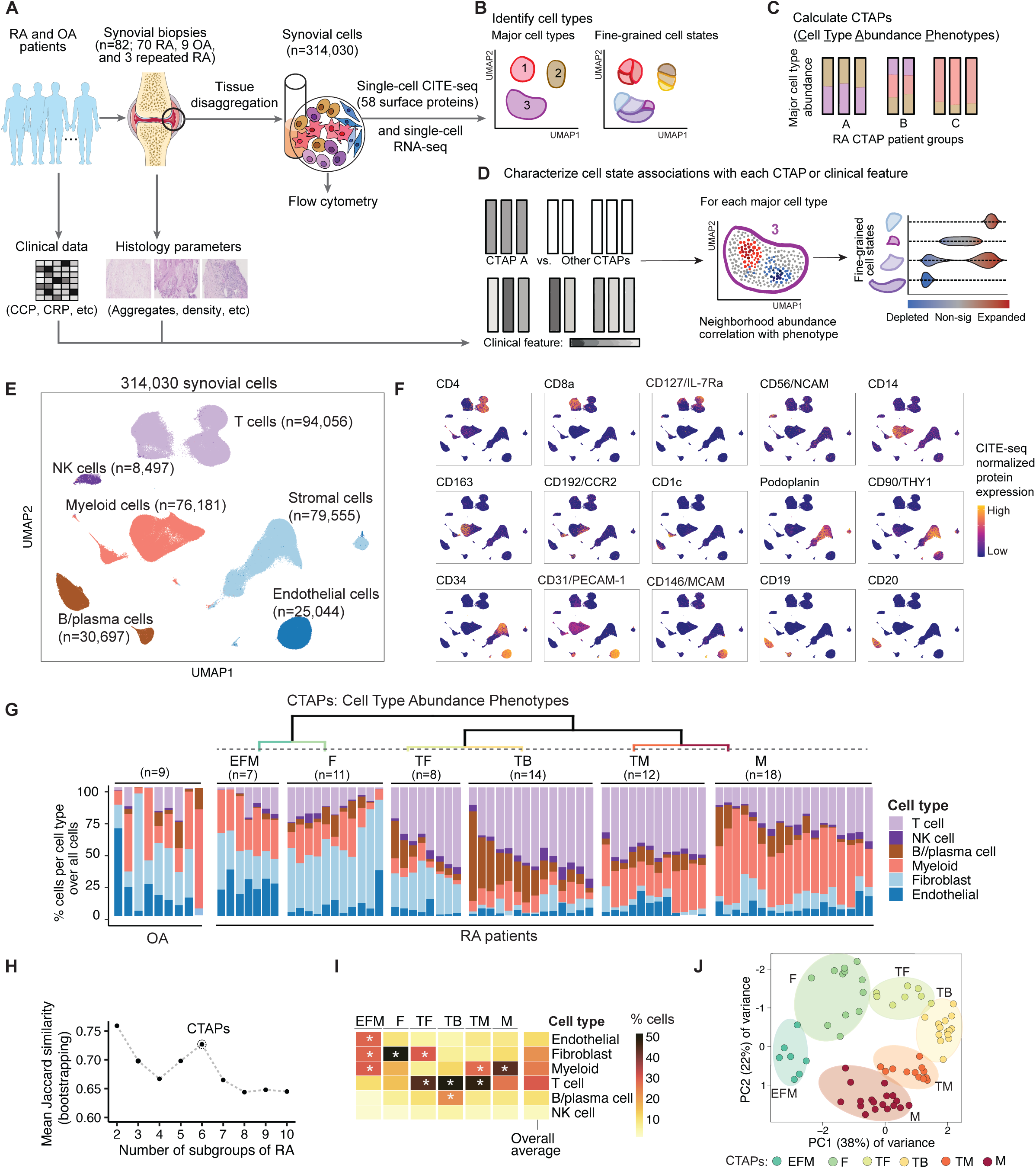
Overview of multimodal single-cell synovial tissue pipeline and cell type abundance analysis reveals distinct RA cell type abundance phenotypes (CTAPs). **A**. Description of patient recruitment, clinical and histologic metrics, synovial sample processing pipeline, and computational analysis strategy, including **B**. identifying major cell types and fine-grained cell states, **C**. definition of distinct RA CTAPs, and **D**. cell neighborhoods associations with each CTAP or with clinical or histologic parameters for each major cell type, **E**. Integrative UMAP based on mRNA and protein discriminated major cell types, **F**. UMAPs of CITE-seq antibody-based expression of cell type lineage protein markers. Cells are colored based on expression from blue (low) to yellow (high), **G**. Hierarchical clustering of cell type abundances captures six RA subgroups, referred to as cell type abundance phenotypes (CTAPs). The nine OA samples are shown as a comparison. Each bar represents one synovial sample, colored by the proportion of each major cell type, **H**. Mean Jaccard similarity coefficient to test CTAP stability by bootstrapping 10,000 times for each tested number of patient subgroups ranging from 2 to 10, **I**. Average proportions of each major cell type among samples in each CTAP. Overall average proportions across all the samples are shown as a comparator. Asterisk represents the proportion that is greater than the overall average for that cell type, **J**. PCA of major cell type abundances. Each dot represents a sample, plotted based on its PC1 and PC2 projections and colored by CTAPs.

### Collection of synovial samples from RA patients

We recruited patients exhibiting moderate to high disease activity (100% with CDAI≥10; 80.6% with DAS28-CRP3≥3.2) and obtained synovial tissue biopsies. To capture the full diversity of RA, we recruited treatment-naive patients (n=28) early in their disease course (mean 2.64 years), methotrexate-inadequate (MTX) responders (n=27), and anti-TNF agent inadequate responders (n=15). The patients were similar in age, sex, disease activity, and other clinical parameters across the three treatment groups (**Supplementary Table 1**). For comparison, we obtained tissues from patients with osteoarthritis (OA, n=9). We assayed a total of 82 synovial tissue samples, including three pairs of samples from RA patients biopsied at two separate times. Three pathologists independently scored each sample for lining layer hyperplasia, cell density, and aggregates^35^, and observed that only cell density was different among patient groups (*p*=0.005, **Supplementary Table 1**).

### Multimodal single-cell integration defines major cell types

We used CITE-seq to simultaneously characterize the full transcriptome and surface expression of 58 proteins, for which we developed and optimized an oligo-conjugated antibody CITE-seq panel spanning key immune and stromal cell lineage and functional markers (**Supplementary Table 2**). We titrated 58 oligo-conjugated antibodies to maximize signal-to-noise (**Methods**). After disaggregating synovial tissue samples, we sorted viable cells for sequencing. A total of 314,030 cells (∼3,800 per sample) passed stringent RNA QC, protein QC, and doublet detection. We also excluded cells with inconsistent cell-type identities based on protein and mRNA (**Supplementary Figure 1A-G**, **Methods**). The proportion of cells within 15 lineage gates in CITE-seq and in flow cytometry correlated across samples (median Pearson r=0.88, **Supplementary Figure 1G-H, Supplementary Table 3**). We integrated surface marker and RNA data using canonical correlation analysis (CCA), corrected batch effects with Harmony^36^, and defined six major cell types: T, B/plasma, natural killer (NK), myeloid, stromal, and endothelial cells (**Figure 1E-F**, **Supplementary Figure 2A-G**, **Methods**).

### Clustering samples on major cell-type abundance to define CTAPs

We quantified the frequency of the six major cell types in each synovial tissue sample (**Figure 1G**). We used these six major cell types instead of finer-grained cell states to create a broad categorization scheme that generalizes easily to many technologies (e.g. flow cytometry) for wide clinical use. We then used hierarchical clustering to classify the spectrum of patient samples into six different synovial cell-type abundance phenotypes (CTAPs). We arrived at six groups because they demonstrated robust in-group similarity with bootstrapping and revealed biological heterogeneity (**Figure 1G-H, Supplementary Figure 2H**, Jaccard index=0.727). We named CTAPs based on dominant cell type(s): 1) endothelial, fibroblast, and myeloid cells (EFM), 2) fibroblasts (F), 3) T cells and fibroblasts (TF), 4) T and B cells (TB), 5) T and myeloid cells (TM), and 6) myeloid cells (M) (**Figure 1I, Supplementary Table 4, Methods**). CTAPs reflect a spectrum of cell-type abundances apparent in principal component analysis (PCA) of cell-type frequencies (**Figure 1J**).

### Characterizing a comprehensive RA synovial cell state atlas

We defined finer-grained cell states and quantified sample abundances within cell types (**Figure 2**). Surface proteins were informative for cell-state delineation in T and B cells (**Supplementary Figure 3A-C**), so we clustered cells on CCA canonical variates (CVs) capturing both RNA and protein data (**Supplementary Figure 3D-F, Supplementary Figure 4, Methods**). For other cell types, proteins were less informative, so we defined clusters from mRNA alone. In total we defined 77 cell states: 24 T cell clusters (n=94,056 cells), 9 B/plasma cell clusters (n=30,697), 14 NK clusters (n=8,497), 15 myeloid clusters (n=76,181), 5 endothelial clusters (n=25,044), and 10 stromal clusters (n=79,555) (**Figure 2A**). Using Symphony^37^, we mapped cell states from our prior study of 5,000 synovial cells^21^ onto these fine clusters^21^; coarse cell states previously identified as associated with RA versus OA were also associated in this data set (**Supplementary Figure 5**, **Supplementary Table 5**).

**Figure 2.**
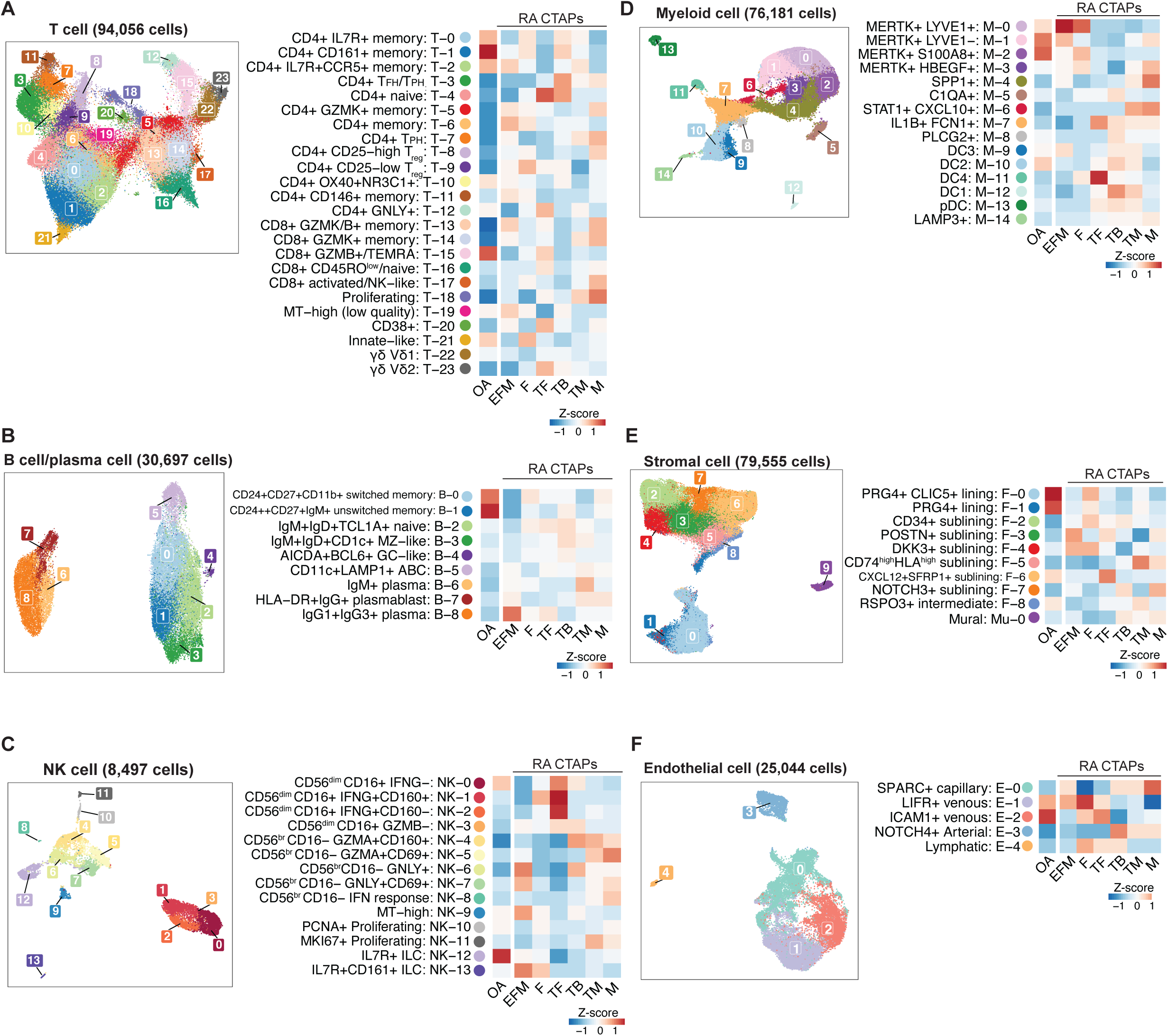
Cell-type-specific single-cell analysis captures 77 distinct cell states in RA synovium. **A-F**. Six cell-type-specific reference UMAPs colored by fine-grained cell state clusters. For each cell type, the heatmap shows the average proportions of each cluster across patient samples in each RA CTAP and OA, scaled within each cluster.

The 24 T cell clusters spanned innate-like states and CD4+ and CD8+ adaptive lineages (**Figure 2A**, **Supplementary Figure 6A-C**). These included states implicated in autoimmunity, such as regulatory CD4^+^ T cells (T_reg_; T-8 and T-9) and T_PH_ and T_FH_ cells (T-3, T-7)^24,28–30,38–41^. T-3 and T-7 both expressed B cell-helper factors CXCL13 and *IL21*. T-7 comprised exclusively T_PH_ cells and expressed more *ICOS*, *IFNG,* and *GZMA*, while T-3 contained T_PH_ and T_FH_ cells expressing the lymphoid homing marker *CCR7* (**Supplementary Figure 6A-D**). T_PH_ cells are known to be expanded in RA compared to OA^21,24^. CD8^+^ subsets expressed different combinations of *GZMB* and *GZMK* (T-13, T-14, T-15), reflecting differential cytotoxic potential. With surface protein data we resolved T cell clusters that were not observed in our earlier study^21^. This included *GNLY*^+^CD4^+^ (T-12), two double-negative (CD4-CD8-) gamma-delta T cell clusters expressing *TRDC* (T-22 and T-23), and a cluster containing double-negative and CD8+ T cells expressing *ZBTB16* (PLZF) that resemble innate-like T cells such as natural killer T cells and mucosal-associated innate T (MAIT) cells (T-21).

We found distinct separation between CD20^+^ (*MS4A1^+^*) B cells and CD138^+^ (*SDC1^+^*) plasma cells (**Figure 2B, Supplementary Figure 7A-D**). CD20+ B cells comprised six clusters, including *IGHM*^+^*IGHD*^+^*TCL1a*^+^ naive (B-2) and two CD27^+^ memory B cell clusters: CD24^+^CD27^+^CD11b^+^ switched memory B cells (B-0) and *CD24^++^CD27^+^IGHM^+^* unswitched memory B cells (B-1). We also identified *CD11C*^+^*CXCR5*^low^ ABCs (B-5)^42–44^, previously noted to be associated with RA relative to OA^21^. B-5 expresses LAMP1, a lysosomal-associated membrane protein that may play a role in B cell antigen-presentation^45^. Additional B-5 genes suggest ABC antigen-presentation^46^ including *HLA-DR* and *CIITA*^47^. We unexpectedly observed CD1c^+^ B cells (B-3) with *CD27* and *IGHD* expression consistent with recirculating extrasplenic marginal zone (MZ) B cells^48–51^. CD1c^+^ MZ-like B cells (B-3) and other non-plasma B cells were high producers of *IL6* and *TNF* (**Supplementary Figure 7D**). We identified *AICDA*^+^*BCL6*^+^ GC-like B cells (B-4) consistent with ectopic germinal center (GC) formation in the synovium^52,53^. Plasma cells were surprisingly diverse and included *HLA-DRA*^+^*MKI67*^+^ plasmablasts (B-7), *IGHM*+ plasma cells (B-6), and more mature *IGHG1*^+^*IGHG3*^+^ plasma cells (B-8). Plasma cell heterogeneity may reflect both *in situ* generation and circulation from the periphery.

We also captured innate lymphocytes, including CD56^br^CD16-NK (8 clusters), CD56^dim^CD16^+^ NK (4 clusters), and CD56^dim^CD16-IL7R+ innate lymphoid cells (ILCs, 2 clusters) (**Figure 2C**, **Supplementary Figure 8A-C**). CD56^br^CD16^-^ NK cells were more abundant (mean 47.6% per donor) than CD56^dim^CD16^+^ NK cells (35.7%) and ILCs (12.9%), consistent with previous observations in gut and lymph nodes^54^. CD56^br^CD16^-^ NK clusters were the only innate lymphocytes expressing *GZMK*, and they variably expressed other genes encoding cytotoxic molecules such as *GZMB* and *GNLY*. CD56^dim^CD16^+^ NK cells had universally high expression of *GZMB, GNLY,* and *PRF1*. *IFNG* was expressed highly in two CD56^dim^CD16^+^ clusters (NK-1, and NK-2) but was also expressed in NK-5 and NK-10. Some activating and inhibitory NK cell receptors were differentially expressed, including *KLRK1* (NKG2D), predominantly expressed by CD56^br^CD16^-^ cells, and *KLRF1* (NKp80) and *FCRL6,* predominantly expressed by CD56^dim^CD16^+^ cells (**Supplementary Figure 8D**). We identified ILCs based on absence of CD56 and CD16 and high expression of CD127 (IL-7Ra) protein^55^. The larger ILC cluster resembled group 3 ILCs (*RORC^+^* NK-12), the functional analog of T_H_17 T cells^55,56^. The smaller CD161+ population resembled group 2 ILCs (*GATA3^+^* NK-12)^55–57^, analogous to T_H_2. We did not see a discrete cluster of T_H_1-analogous group 1 ILCs, which may have co-clustered with NK cells.

We identified 15 myeloid clusters spanning tissue macrophages, infiltrating monocytes, conventional and plasmacytoid dendritic cells (**Figure 2D**). CD68 and CCR2 protein expression discriminate tissue macrophages from infiltrating monocytes (**Supplementary Figure 9A-C**). Three tissue macrophage clusters (M-0, M-1, M-2) in RA synovium were also found at high frequencies in OA synovium and display a phagocytic phenotype with high expression of CD206 (*FOLR2*), CD163, *MERTK* and *MARCO* (**Supplementary Figure 9B,D**), suggesting homeostatic debris-clearing function^58,59^. *LYVE1* expression on tissue macrophages (M-0) may indicate a perivascular function^25,60^. Infiltrating monocytes included a sizable *IL1B^+^FCN1^+^HBEGF^+^* pro-inflammatory subset (M-7), likely derived from classical CD14^high^ monocytes, which we previously described^21,25^. A *STAT1^+^CXCL10^+^* subset (M-6) likely derives from non-classical CD14^low^CD16^high^ monocytes and expresses interferon-response gene signatures; these cells are enriched in the inflamed lung from COVID-19 pneumonia, colon from Crohn’s disease, and tumors^23,61,62^. *MERTK*^+^*HBEGF*^+^ (M-3) and *SPP1*^+^ (M-4) bridged infiltrating monocytes and tissue macrophages; both expressed high levels of *SPP1*, a marker of bone-marrow-derived macrophages^63,64^ suggesting a transition from an inflammatory monocyte to a more phagocytic phenotype of tissue macrophages. We identified four DC populations corresponding to subsets described by Villani *et al*^65^. Reflecting their respective antigen presentation capacities, DC1 (M-12) expressing *CLEC9A* and *THBD* (CD141) cross-present extracellular antigens to CD8 T cells, while DC2 and DC3 (M-10, 9) are *CLEC10A*^high^ cells that activate and polarize CD4 T cells^65^ (**Supplementary Figure 9D**). DC4 (M-11) expresses genes found in CD14^+^ monocytes such as *IL1B* while also displaying a strong IFN signature. Lastly, we identified a fifth DC subset (M-14) with high expression of endosomal marker *LAMP3*^66^.

In the stroma, fibroblasts were divided broadly into lining (*PRG4*^high^) and sublining (*THY1*^+^ *PRG4*^low^) (**Figure 2E, Supplementary Figure 10A-F)**. As previously described, lining fibroblasts (F-0, F-1) were relatively depleted in RA and enriched in OA synovium, while sublining fibroblasts separated into *HLA-DRA^+^*, *CD34^+^*, and *DKK3^+^* groups^21,67,68^ (**Supplementary Table 6**). Lining fibroblasts subdivided into *PRG4^+^CLIC5^+^* (F-0), *PRG4^+^* (F-1), and an *RSPO3^+^* population (F-8) with an intermediate lining/sublining phenotype. The *CD34^+^* sublining fibroblast cluster (F-2) highly expressed *PI16* and *DPP4* (CD26), suggesting they may be fibroblast progenitors^69^. *CXCL12^+^* fibroblasts included an inflammatory *CD74*^high^*HLA*^high^ cluster (F-5) with high HLA expression, and a *CXCL12^+^SFRP1^+^* cluster (F-6) with the highest levels of *IL6,* a proven drug target in RA^70–72^. The inflammatory signature in F-5 and F-6 suggest an inflammatory phenotype driven by cytokine activation by infiltrating immune cells^73^. The stromal compartment also included a small cluster of *NOTCH3^+^MCAM* (CD146)^+^ mural cells (Mu-0).

Endothelial cells separated into *NOTCH4^+^* arteriolar (E-3), *SPARC^+^* capillary (E-0), *CLU^+^* venular (E-1, E-2), and *LYVE1^+^PROX1^+^* lymphatic endothelial cells (LEC, E-4) (**Figure 2F**, **Supplementary Figure 10G-K**). The majority (53%) were venular and further subdivided into *LIFR^+^* (E-1) and *ICAM1^+^* (E-2); these cells had high expression of inflammatory genes such as *IL6* and *HLA*, along with genes that facilitate the transmigration of leukocytes into tissue such as *ICAM1* and *SELE* (E-selectin)(**Supplementary Figure 10I**)^74^. Arteriolar cells expressed high levels of *CXCL12, LTBP4, NOTCH4,* and NOTCH ligand *DLL4*. *SPARC^+^* capillary cells expressed collagen and extracellular matrix genes. LECs represented a small number of cells (n=324) with high expression of *CCL21* and *FLT4*^75,76^.

For each sample, we calculated the proportion of each cell cluster within each cell type. Then, we calculated the average of these cluster proportions within each RA CTAP and OA (**Figure 2**). These values are independent of cell-type abundance differences since they are calculated relative to each cell type. For example, these values may reflect the relative abundance of *IL1B^+^* macrophages among all myeloid cells, regardless of the total number of myeloid cells in a sample. We observed reported differences in RA compared to OA, including an expansion of sublining fibroblasts relative to lining fibroblasts, and expansion of *IL1B^+^* macrophages relative to *MERTK^+^* macrophages.

### CTAPs are characterized by specific cell states

We next set out to quantify how the composition of fine-grained cell states differed between CTAPs. To accurately identify cell-states associated with individual CTAPs within each given cell type, we used co-varying neighborhood analysis (CNA)^77^. CNA tests highly granular “neighborhoods”—small groups of phenotypically similar cells—rather than larger clusters and accounts for age, sex, and cell count per sample. CNA associations suggest that certain single-cell-resolution states within each cell type are more likely to be found in samples from one CTAP than others. After identifying CTAP-associated neighborhoods, we defined the canonical cell states that contain those neighborhoods to infer biologic meaning. In these analyses, we use “expanded” and “depleted” to refer to changes in relative abundance within a cell type, but notably these changes may not reflect a change in absolute cell numbers relative to total number of cells.

We observed skewed T cell neighborhoods in CTAP-TB (permutation *p*=0.046) (**Methods**, **Figure 3A-B, Supplementary Figure 6E**, **Supplementary Table 6**). T cell neighborhoods among CD4^+^ T_FH_/T_PH_ (T-3) and CD4+ T_PH_ (T-7) cells were expanded, while T cell neighborhoods among cytotoxic CD4^+^*GNLY^+^* (T-12) and CD8^+^*GZMB^+^* cells (T-15) were depleted. Recognizing that T_FH_ and T_PH_ cells differentiate B cells towards antibody production^24,78^, we tested B cells for association to CTAP-TB (permutation *p*=0.03). We observed expanded neighborhoods in memory B (B-0 and B-1) and ABC (B-5) clusters, while IgG1^+^IgG3^+^ and IgM^+^ plasma cells (B-8, B-6) were relatively depleted (**Figure 3C-D, Supplementary Figure 7E**, **Supplementary Table 6**). We note that though plasma cells are depleted among B/plasma cells in CTAP-TB, B and plasma cells overall are enriched among total cells in CTAP-TB (23% compared to 1-10% in other CTAPs) (**Figure 1I, Supplementary Table 4**). While T_PH_ (T-7), T_FH_/T_PH_ (T-3), and ABC (B-5) cells are enriched in CTAP-TB, they are present in all six CTAPs (**Supplementary Figures 6E and 7G, Supplementary Table 6**). In contrast, GC cells (B-4) were almost exclusively found in CTAP-TB (**Supplementary Figure 7G**). Consistent with a role for T_FH_/T_PH_ and IL21 in ABC generation^43^ and plasma cell differentiation, the frequency of ABCs (B-5) amongst B/plasma cells correlated with the proportion of T_PH_ (T-7) (Pearson r=0.50, *p*=3.7e-6, **Figure 3E**) and T_FH_/T_PH_ (T-3) amongst T cells (Pearson r=0.24, *p*=0.034, **Supplementary Figure 7F**).

**Figure 3.**
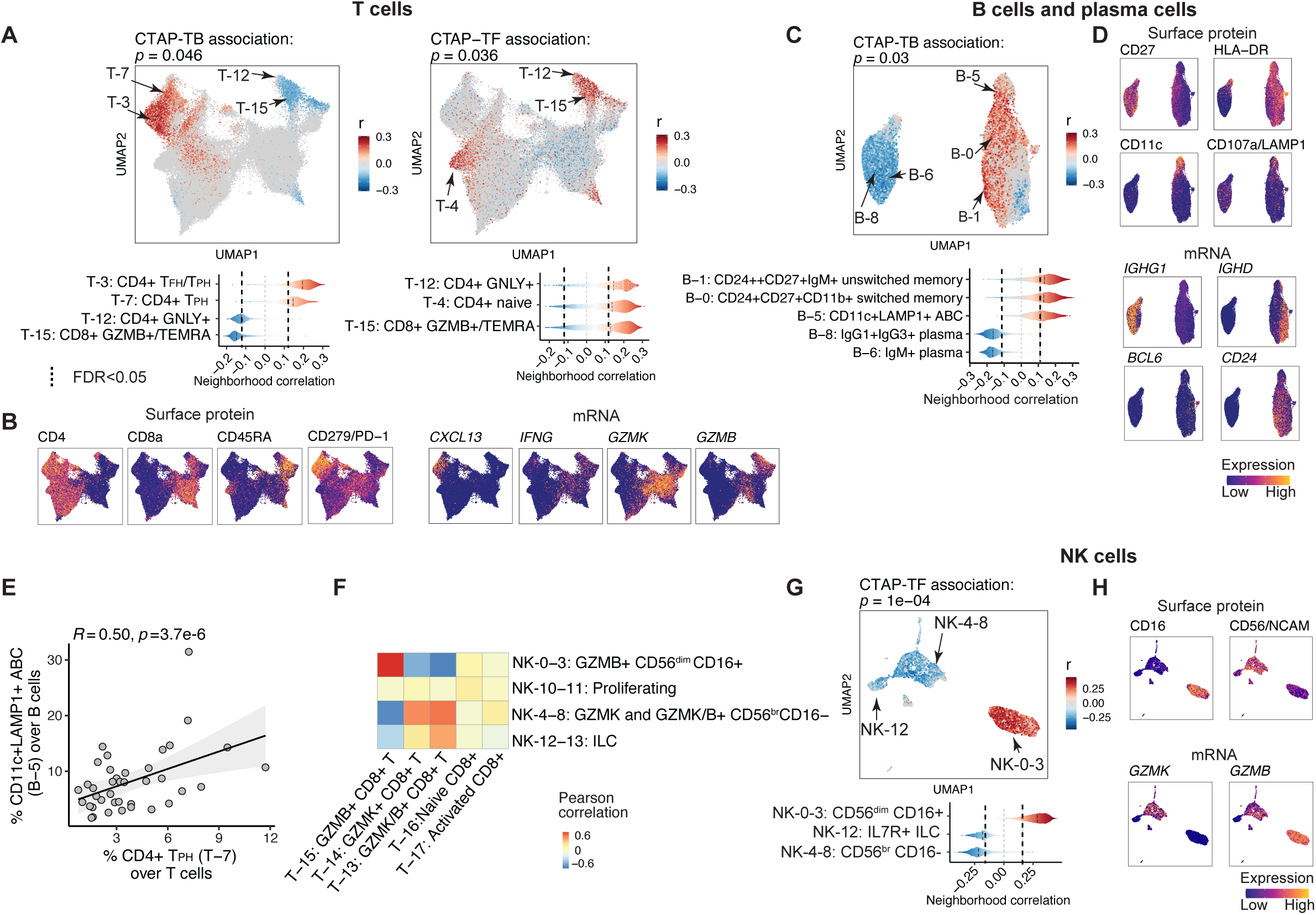
Different T cell, B cell, and NK cell populations are associated with RA CTAPs. **A**, Associations of T cell neighborhoods with CTAP-TB and CTAP-TF. P-values are from the CNA test for each CTAP within T cells. For all CNA results, cells in UMAP are colored in red (positive) or blue (negative) if their neighborhood is significantly associated with the CTAP (FDR < 0.05), and gray otherwise. Distributions of neighborhood correlations are shown for clusters with >50% of neighborhoods correlated with the CTAP at FDR>0.05, **B**. Expression of selected surface proteins and transcripts among T cells. For all expression UMAPs, cells are colored from blue (low) to yellow (high), **C**. Associations of B/plasma cell neighborhoods with CTAP-TB, **D**. Expression of selected surface proteins and transcripts among B/plasma cells, **E**. Percentage of T_PH_ (T-7) out of T cells and CD11c+ LAMP1+ ABCs (B-5) out of B/plasma cells for each donor sample, represented by points. R and p-value are calculated from Pearson correlation, **F**. Heatmap colored by Pearson correlation between per-donor CD8 T cell and NK cell cluster abundances, **G**. Associations of NK cell neighborhoods with CTAP-TF. **H**. Expression of selected surface proteins and transcripts in NK cells.

T cell neighborhoods enriched in CTAP-TF (permutation *p*=0.036) mainly consisted of cytotoxic CD4^+^*GNLY^+^* (T-12) and CD8^+^*GZMB^+^* cells (T-15) (**Figure 3A, Supplementary Figure 6E**, **Supplementary Table 6**). Similarly, NK cell neighborhoods were altered in CTAP-TF (permutation *p*=1e-4), and these neighborhoods contained *GZMB*-expressing CD56^dim^CD16^+^ NK cells (NK-0-3) (**Figure 3G-H, Supplementary Figure 8E**). The *GZMB^+^* (NK-0-3) proportion of NK cells correlated with the *GZMB^+^* (T-15) proportion of T cells (Pearson r=0.63, *p*=4.87×10^-10^, **Figure 3F**). This suggests that a subset of RA samples is enriched in *GZMB^+^* NK and T cells expressing high *IFNG* (**Supplementary Figure 6D**, **Supplementary Figure 7D**). Conversely, we observed that CD8^+^ T cells expressing *GZMK* (T-13/14) correlated with NK cells expressing *GZMK* (NK-4-8, Pearson r=0.51, *p*=1.41×10^-6^, **Figure 3F**), suggesting that *GZMK*-expressing CD8 T and NK cells share a transcriptional program that may result from their tissue environments.

CTAP-TF also exhibited specific expansions of fibroblast subpopulations (permutation *p*=0.048, **Figure 4A-B**). Specifically, *CXCL12^+^SFRP1^+^* sublining fibroblasts (F-6) were disproportionately expanded in CTAP-TF. These *CXCL12^+^SFRP1^+^* sublining fibroblasts highly expressed *IL6* but did not express HLA-DR genes.

**Figure 4.**
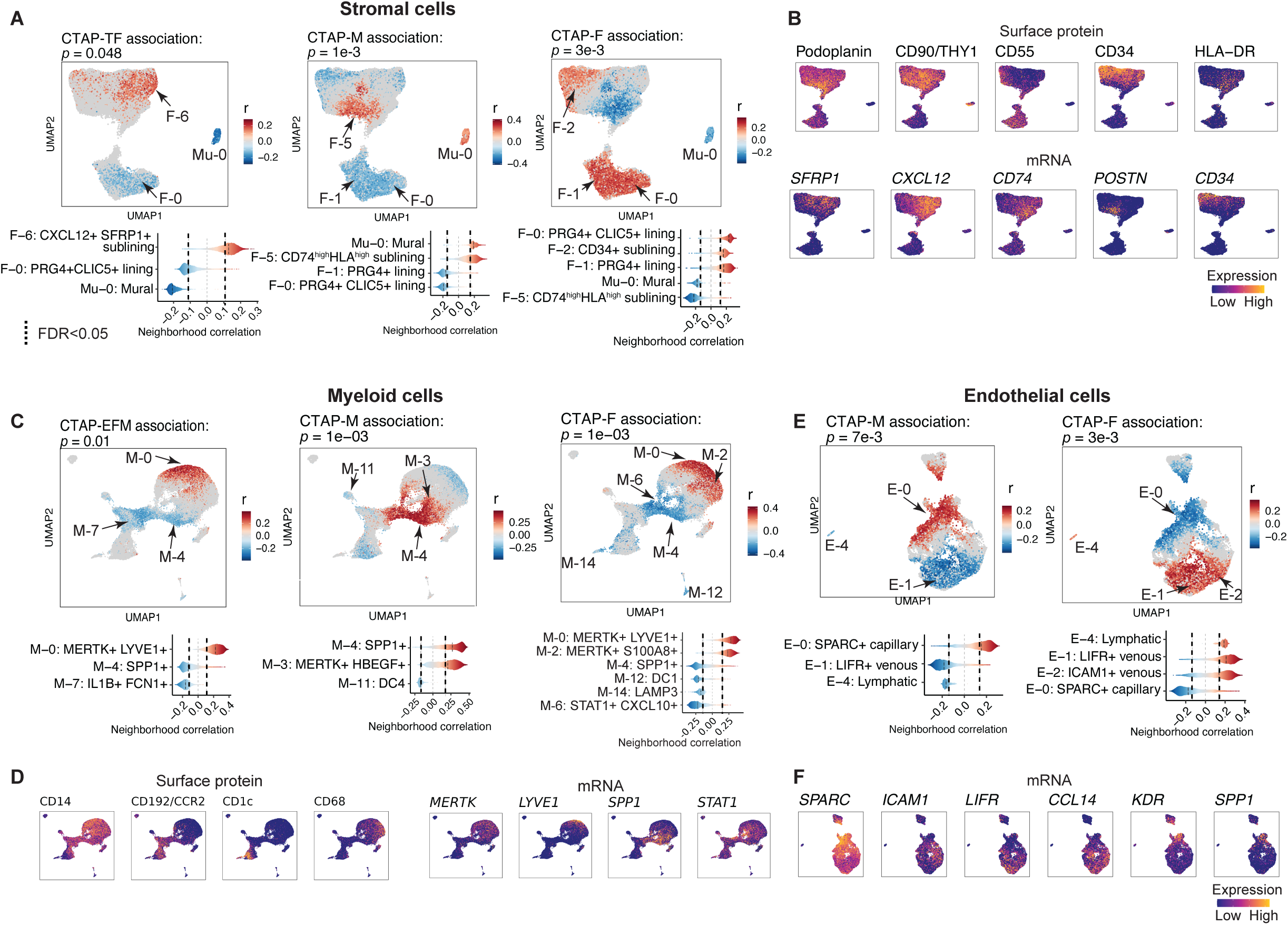
Different stromal, myeloid, endothelial cell populations are associated with RA CTAPs. **A**. Association of stromal cell neighborhoods with CTAP-TF, CTAP-M, and CTAP-F. For all CNA results, cells in UMAPs are colored in red (positive) or blue (negative) if their neighborhood is significantly associated with the CTAP (FDR < 0.05), and gray otherwise. Distributions of neighborhood correlations are shown for clusters with >50% of neighborhoods correlated with the CTAP at FDR>0.05, **B.** Expression of selected surface proteins and transcripts among stromal cells. For all expression UMAPs, cells are colored from blue (low) to yellow (high), **C**. Association of myeloid cell neighborhoods with CTAP-EFM, CTAP-M, and CTAP-F, **D.** Expression of selected surface proteins and transcripts among myeloid cells, **E**. Association of endothelial cell neighborhoods with CTAP-M and CTAP-F, **F.** Expression of selected surface proteins and transcripts among endothelial cells.

Myeloid populations were different in CTAP-M compared to other CTAPs (permutation *p*=1e-3). Cell neighborhoods within *SPP1^+^* (M-4) and *MERTK^+^HBEGF^+^* (M-3) bone marrow-derived macrophages were enriched in CTAP-M suggesting recruitment of inflammatory monocytes and transition to macrophage function (**Figure 4C-D**). Furthermore, in CTAP-M, *CD74*^high^*HLA*^high^ sublining fibroblast neighborhoods (F-5) were expanded relative to stromal cells (permutation *p*=1e-3) and *SPARC*^+^ capillary cells (E-0) were expanded relative to endothelial cells (permutation *p*=7e-3, **Figure 4A-B, E-F**). Interestingly, the neighborhoods expanded in CTAP-M were depleted in CTAP-F, while neighborhoods depleted in CTAP-M were enriched in CTAP-F. Specifically, subpopulations like lining (F-0 and F-1) and CD34^+^ sublining (F-2) fibroblasts (permutation *p*=3e-3), *MERTK^+^LYVE1^+^* (M-0) and *MERTK^+^S100A8^+^* (M-2) macrophages (permutation *p*=1e-3), and *LIFR^+^* venular (E-1) and *ICAM1^+^* venular (E-2) endothelial cells were expanded in CTAP-F (permutation *p* = 3e-3) and depleted in CTAP-M. Notably, the pro-inflammatory *IL1B^+^* macrophages^21^ (M-7), known to be expanded in RA patients in general^21^, were lower in frequency in CTAP-EFM relative to other CTAPs (**Figure 4C**).

### Cell states and CTAPs associated with histology and clinical metrics

In addition to association with CTAPs (**Figure 5A**), cell neighborhoods may also be associated with histologic features of RA synovium, which are useful in clinical practice and reflect disease pathogenesis^79–81^. Using CNA, we identified transcriptional neighborhoods associated with histology, accounting for age and sex (**Methods**). We scored samples for Krenn histologic inflammation and lining layer domains, in addition to discrete histologic cell density and aggregate abundance, reflecting inflammatory cell infiltration and organization respectively (**Supplementary Figure 11A**). T cells were associated with aggregate scores (permutation *p*=0.0088), driven by expanded T cell neighborhoods in CD4^+^ T_FH_/T_PH_ (T-3), consistent with their role in organizing secondary lymphoid structures^82,83^ (**Supplementary Figure 11B,** **Figure 5A**). IgM+ plasma cells (B-6), plasmablasts (B-7), and ABCs (B-5) were also positively associated with aggregates (permutation *p*=0.007) (**Supplementary Figure 11B,** **Figure 5A**). In similar analysis of NK cell neighborhoods, CD56^br^CD16^-^GZMA^+^CD160^+^ cells (NK-4) were positively associated with density and aggregate scores (permutation *p*=3e-04 and 1e-04, respectively) (**Supplementary Figure 11B**); this population also contained cell neighborhoods relatively enriched in CTAP-TB (**Figure 2**), although the functional role of these cells in follicle-rich synovium is less clear. Inflammatory myeloid neighborhoods within *STAT1^+^CXCL10^+^* (M-6), *SPP1^+^* (M-4) and inflammatory DC3 (M-9) (**Supplementary Figure 11B**, **Figure 5A**) were associated with density (permutation *p*=0.005).

**Figure 5.**
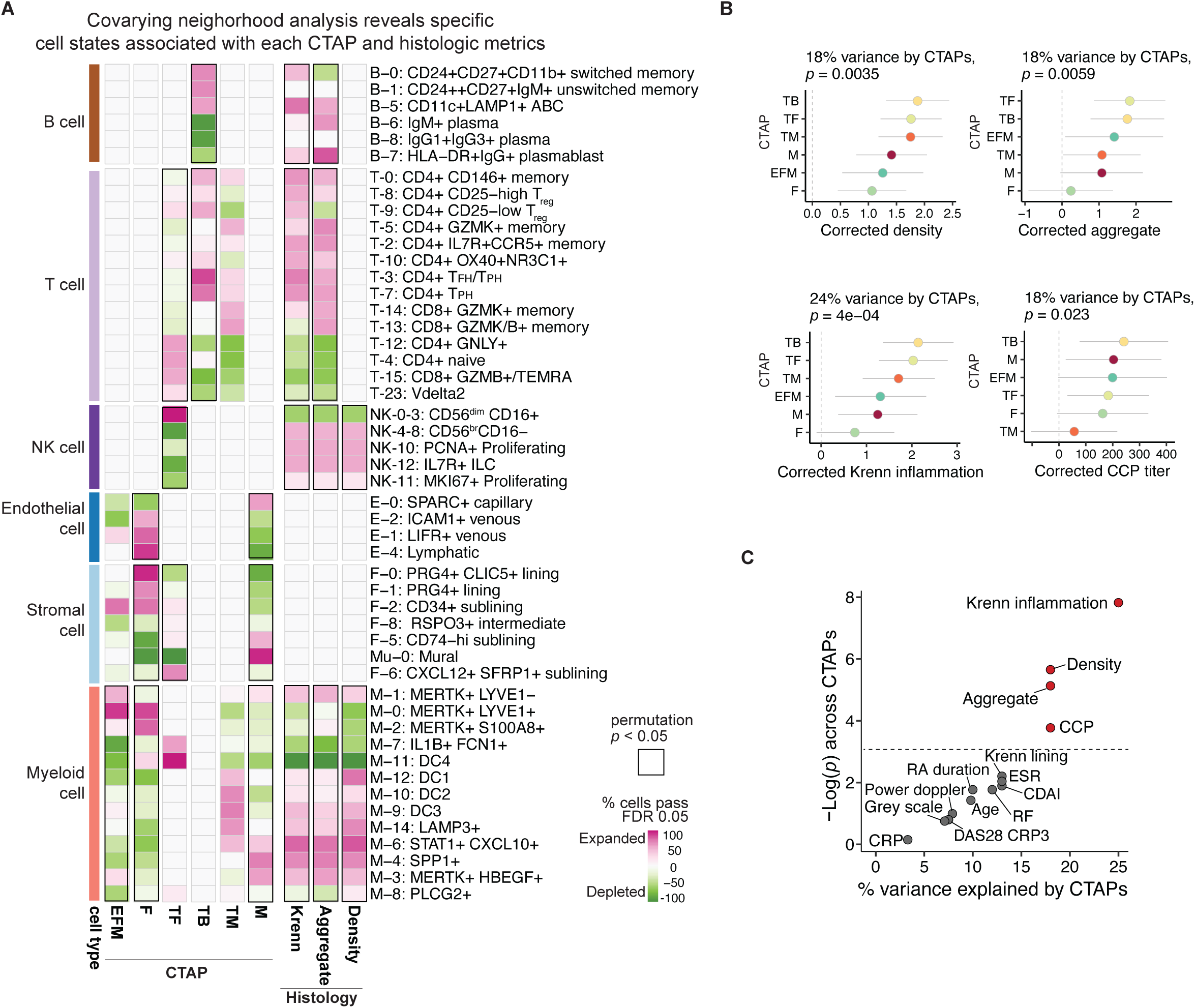
Single-cell covarying neighborhood analysis reveals significant association of cell states with disease indicators. **A**. Heatmap of CNA associations of specific cell states with each RA CTAP. Colors represent % cell neighborhoods from each cell state with local (neighborhood-level) phenotype correlations passing FDR < 0.05 significance from white to pink (expanded) or green (depleted). Cell types significantly associated globally (cell-type-level) with a phenotype at permutation *p* < 0.05 are boxed in black, **B**. Association between clinical features and CTAPs, adjusting covariates for age, sex, cell number, and clinical collection site. Percentage of variance explained by CTAPs alone and p-value are calculated with ANOVA tests. 95% confidence intervals are shown. **C**. Clinical, demographic, and histologic metrics plotted by percentage of variance explained by CTAPs and the ANOVA p-value for its association with CTAPs. Features in red are significant at ANOVA *p* < 0.05.

We wanted to understand if histologic and clinical measures are explained by CTAPs, taking age, sex, cell count, and clinical collection site into account (**Methods**). CTAPs account for 18% variance of histologic density (*p*=0.0035) and 18% of variance for aggregates (*p*=0.0059), with CTAP-TB and CTAP-TF having the highest scores for both (**Figure 5B**, **Supplementary Figure 12A**). Consistent with these observations, CTAPs are associated with Krenn inflammation scores (*p*=4e-04), but not with Krenn lining scores (*p*=0.11) (**Figure 5B**, **Supplementary Figure 12B**). CTAP-F, CTAP-EFM, and CTAP-M have the lowest scores for all histological parameters (**Figure 5B**).

The presence of CCP autoantibodies and rheumatoid factor subcategorize RA patients as seropositive or seronegative. Patients with positive CCP have more severe disease and radiographic progression^84,85^. CCP titer values differed across CTAPs (*p*=0.023, 18% variance), with CTAP-TB having the highest CCP (mean=292) (**Figure 5B**), even after restricting the analysis to seropositive patients (*p*=0.0047) (**Supplementary Figure 12C**).

Intriguingly, CTAPs were independent of most clinical variables including disease activity, clinical inflammatory markers, smoking history, total swollen joint counts, and sex (**Figure 5C, Supplementary Table 10, Supplementary Figure 12D-L**). CTAPs were also mostly independent of anatomic category and clinical sites (**Supplementary Figure 12H-I**). Patients in CTAP-EFM tended to be older and have longer-standing RA than patients in other CTAPs and were mostly TNFi-inadequate responders (**Supplementary Figure 12J-K**), although these associations were not statistically significant.

### CTAPs feature disease-relevant cytokine profiles

We recognized that cell states differentially expressed specific effector molecules, such as cytokines and their receptors (**Supplementary Figure 13**). Most cytokines and chemokines are produced predominantly by one cell type (**Figure 6A**). For key cytokines produced by multiple cell types, we quantified the relative contributions of each cell type. For example, we found that roughly equal numbers of T cells and myeloid cells express *TNF* while stromal, endothelial, and B cells dominate among *IL-6*-expressing cells (**Figure 6B**).

**Figure 6.**
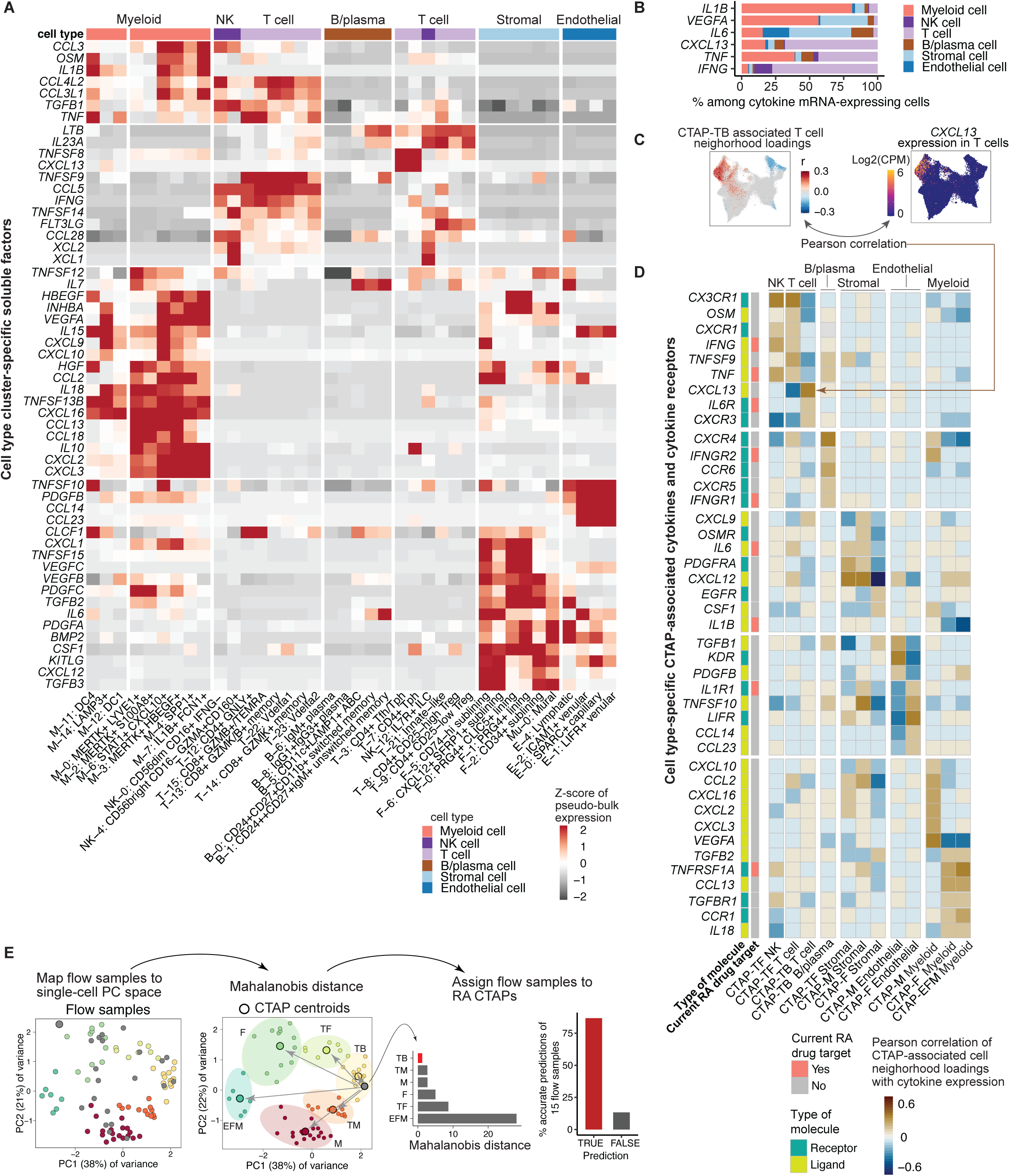
Cell type clusters and CTAPs feature distinct disease-relevant soluble factor and receptor profiles. **A**. Expression profiles of cell type cluster-specific soluble factors, **B**. Percent contribution among cytokine mRNA-expressing cells from each major cell type, **C**. Expression of representative cytokine, *CXCL13*, that is significantly correlated with CTAP-associated cell neighborhoods. Cells in UMAPs of CTAP associations are colored in red (positive) or blue (negative) if their neighborhood is significantly associated with the CTAP (FDR < 0.05), and gray otherwise. Cells in expression UMAPs are colored from blue (low) to yellow (high), **D**. With a heatmap, we visualized the cytokines and receptors whose expressions are significantly correlated (r > 0.5) with CTAP-associated cells; we then hierarchically clustered them based on cell type-specific CTAPs. For each gene, receptor/ligand designation and current RA drug target status are labeled, **E**. Pipeline and results to map and classify flow cytometry samples by single-cell RA CTAPs. Bar plot shows accuracy of flow sample classification (i.e., assigned to the same CTAP as a single-cell sample from the same patient).

Next, we linked these key effector molecules to CTAPs to complement the previous analyses where we identified clusters overlapping with associated cell neighborhoods. To do this, we correlated CTAP neighborhood association scores with expression of key cytokines and receptors to identify soluble factors produced by CTAP-associated cell states.

CTAP-TB T cell neighborhood association scores correlated with expression of T_FH_/T_PH_-marker *CXCL13* in T cells (**Figure 6C-D**), consistent with the observation that associated T cell neighborhoods were in the T_FH_/T_PH_ clusters (**Figure 3A**). In contrast, CTAP-TF-associated T cell neighborhoods were associated with expression of *IFNG* and *TNF*, expressed by cytotoxic (*GZMB*^+^ or *GNLY^+^*) CD8^+^ and CD4^+^ T cell populations (**Figure 3A**, **Figure 6D**). NK cell populations enriched in CTAP-TF also expressed high *IFNG* and *TNF* (**Figure 3G**, **Figure 6D**). These results suggest that *TNF* and *IFNG* may be intrinsic to the molecular environment of CTAP-TF.

Analysis of myeloid cell neighborhoods in CTAP-EFM, CTAP-F, and CTAP-M also highlighted key cytokines (**Figure 6D**). CTAP-M myeloid neighborhood association scores correlated with expression of chemokines that related to activity of myeloid cells and neutrophils, *CXCL10* and *CCL2* (**Figure 6D**), and angiogenic factors *CXCL16* and *VEGFA*. In CTAP-M, endothelial cell neighborhood association scores correlated with *KDR* (VEGF receptor 2) (**Figure 6D**), consistent with the prevalence of capillary cells in CTAP-M^86^. In contrast, in CTAP-F, *LIFR*^+^ and *ICAM1*^+^ venous endothelial cells expressed high levels of *CCL14,* whose cognate receptor *CCR1* was highly expressed by *MERTK*^+^ macrophages, offering a potential mechanism for the enrichment of this macrophage subset (**Figure 4C-D**, **Figure 6D**).

### CTAPs serve as a reference to map data from other patients and cohorts

Our study included three patients with replicate biopsies obtained from the same joint (98, 105, and 190 days) after the initial biopsy. We assessed the stability of CTAP phenotypes over time between repeated and baseline samples. We found that the cell-type composition of repeat biopsies was similar to the initial biopsy (mean Mahalanobis distance=1.55, permutation *p*=0.073) (**Supplementary Figure 14A-B**), though a larger study would be necessary to understand how dynamic CTAPs are in a given patient.

Given the potential benefits of categorizing synovial tissues from future RA studies into CTAPs, we next examined whether samples can be classified into CTAPs using a lower-resolution technology such as flow cytometry. We built a Mahalanobis-distance-based nearest-neighbor classifier, and we were able to accurately replicate CITE-seq-based CTAP assignments based on flow cytometry data (accuracy=87%, **Figure 6E**, **Supplementary Figure 14C-D, Methods**). Since CTAPs appear to correlate with known drug targets (**Figure 6D**) and can be assigned even with flow cytometry, we expect that CTAPs can be used to systematically query RA heterogeneity across technologies to improve the granularity of clinical studies and trials and potentially to guide therapy selection.

## Discussion

We constructed a comprehensive synovial tissue inflammation reference of >314,000 single cells. This clinically phenotyped RA atlas can be used to classify single-cell data from other RA patients, identify shared pathways across diseases, and identify novel drug targets. We observed that inflamed tissue samples from RA patients have diverse cellular composition that is captured in six CTAPs.

CTAPs represent categories of RA characterized by the presence of certain cell states and the absence of others. We observed that some previously identified pathogenic cell states in RA are expanded in specific CTAPs. For example, CD4+ T_FH_ and T_PH_ cells, generally observed to be enriched among T cells in RA compared to OA^24^, are present in all CTAPs but are most expanded in CTAP-TB. These T cell states are enriched along with ABCs and memory B cells, consistent with the formation of B and T cell aggregates. Independent work has shown that B cell activation pathways are active in the setting of human autoimmunity^44^. Importantly, prior work has focused on peripheral blood and normal secondary lymphoid tissue^87–89^, and it remains unknown how this translates to B cell activation and ectopic lymphoid reactions in RA synovium. Our work suggests the presence of two synovial B cell activation pathways, including conventional germinal center responses and extra-follicular pathways, the latter characterized by CXCR5-ABCs and T_PH_. The rarity of GC dark-zone B cells and abundance of ABCs suggest the prominence of extra-follicular activation pathways in RA synovium. Other novel findings from the B-cell analyses include genes associated with antigen presentation in the ABCs, the presence of CD1c-MZ-like B cells previously described in Sjogren’s disease salivary glands^90^, and the heterogeneity of plasma cells.

In other cell types as well, CTAPs delineate RA subsets where established cell states of interest may be more or less prominent. For example, among fibroblasts, prior studies have found that inflammatory sublining cells expressing *HLA-DR*, *CXCL9*, *CXCL12*, and *IL6* are known to be enriched in RA compared to OA^21,68^. Here, we find that these inflammatory sublining cells are composed of several subpopulations—some of which, specifically *CXCL12^+^* and *CD74*^high^*HLA*^high^ cells, were enriched in CTAP-TF and CTAP-M, respectively. These findings may reflect different axes of inflammatory fibroblast phenotypes, likely involving signals exchanged with surrounding leukocytes. Interestingly, CTAP-M, where *CD74*^high^*HLA*^high^ fibroblasts are enriched, also exhibits specific enrichment of *MERTK^+^HBEGF^+^* and *SPP1+* (osteopontin) macrophages, and several other myeloid populations (e.g. *IL1B*+ *FCN1*+) are also prominent. These and other instances of co-enriched populations (e.g. *GZMK*+ versus *GZMB*+ CD8 T cells and NK cell subsets) inspire new questions about cell-cell interactions underlying inflammatory phenotypes in RA synovium and potentially other tissues and diseases.

CTAPs are associated with histologic and serologic (CCP) parameters but not with drug history and clinical disease activity metrics in this study. We argue that CTAPs from biopsies offer independent information from what physician assessments offer. Our study does not address the evolution of CTAPs in patients over time. We anticipate future longitudinal studies to investigate CTAP changes over time along with treatment effects. In our limited assessment of three patients, we noted minimal evolution of CTAPs despite treatment changes.

Targeting the specific cell subsets enriched in a given CTAP may be key in personalized RA treatment. For example, abrogating T-B cell communication with B cell-depleting antibodies (e.g. rituximab) or blocking costimulation (e.g. abatacept) in CTAP-TB may break the pathogenic mechanisms that drive inflammation in these patients^18,82^. Conversely, patients with CTAP-TF and CTAP-M feature fibroblast populations with high IL-6, an established target of current FDA-approved treatments of RA (e.g. tocilizumab). CTAP-TF and CTAP-M feature abundant *IFNG*-expressing cells or IFN-associated gene signatures, suggesting that these patients may respond effectively to JAK inhibitors (e.g. tofacitinib, upadacitinib). Lastly, other CTAPs, such as CTAP-EFM and CTAP-F, currently have no obvious targets of currently available treatments and warrant further focused study. Thus, CTAPs represent valuable molecular classifications of RA that may drive the search for new treatments.

The CTAP paradigm provides a tissue classification system that captures coarse cell-type and fine cell-state heterogeneity. Importantly, CTAPs use global cell-type frequencies and are thereby an accessible tool to categorize heterogeneity of tissue inflammation using multiple technologies. The model presented here may serve as a powerful prototype to classify other types of tissue inflammation, including in other immune-mediated diseases. A deeper understanding of the heterogeneity of tissue inflammation in RA and other autoimmune diseases may shed new light on disease pathogenesis and reveal new treatment targets.

## Supplementary Figures

**Supplementary Figure 1. Detailed single-cell CITE-seq quality control**. **A**. Quality of the cells based on number of genes detected and percent mitochondrial UMIs (%MT), **B**. Percentage of good quality cells for sample-level QC, **C**. Doublet detection using Scrublet, **D**. UMAP of the number of genes and UMIs detected, **E**. Number of cells remaining after each step of QC, **F**. Distributions of cell type lineage antibody staining from CITE-seq determine percentage of major cell types based on the thresholds (red line) including % CD45^+^ cells, % T cells based on CD4 antibody, % B cells based on CD20^+^, % macrophages based on CD14^+^, % endothelial cells based on CD146^+^, and % fibroblasts based on PDPN^+^, **G.** Representative gating of flow cytometry data to quantify selected synovial cell populations, **H.** Concordance of single-cell CITE-seq antibody staining with an analogous gating schema for flow cytometry. For flow gating, we determined % CD45^+^ based on CD45^+^ over all live cells, % T cells based on CD45^+^CD3^+^ over all live cells, % B cells based on CD45^+^CD3^-^CD14^-^CD20^+^, % macrophages based on CD45^+^CD14^+^, % fibroblasts based on CD45^-^CD146^-^CD31^-^, % endothelial cells based on CD45^-^CD146^+^CD31^+^, % CD4 T cells based on CD45^+^CD3^+^CD4^+^, % HLA^+^ CD4 T cells based on CD45^+^CD3^+^CD4^+^HLA-DR^+^, % CD8 T cells based on CD45^+^CD3^+^CD8^+^, % HLA^+^ CD8 T cells based on CD45^+^CD3^+^CD8^+^HLA-DR^+^, % PD1^+^ CD4 T cells based on CD45^+^CD3^+^CD4^+^PD1^+^, % HLA+ fibroblasts based on CD45^-^CD146^-^CD31^-^HLA^+^, % sublining fibroblast based on CD45^-^ CD146^-^CD31^-^CD90^+^, % CD27^+^ B cells based on CD45^+^CD3^-^CD14^-^CD20^+^CD27^+^, and % CD11c^+^ B cells based on CD45^+^CD3^-^CD14^-^CD20^+^CD11c^+^, respectively.

**Supplementary Figure 2. Single-cell CITE-seq integrative analysis**. **A**. CCA-based pipeline for integrating mRNA and protein expressions, **B**. Concordance between average mRNA expression and the correlations of corresponding protein and mRNA expression. Black line represents the linear best fit line and the shaded region represents the 95% confidence interval, **C.** Sample sources (*n*=82) in the UMAP space, paired with sensitivity analyses of Harmony parameters based on LISI scores to measure mixture levels on **D**. samples and **E**. cell types, **F**. The effect of varying the selected number of antibodies based on each antibody’s specificity: KL divergence equals 0.5 (25 proteins), 0.3 (36 proteins), and 0 (58 proteins), while also varying the number of highly variable genes used: 500/sample (3,164 genes in total) and 1,000/sample (5,751 genes in total) on the mRNA and protein integrative analysis. We used the top 1,000 most variable genes per sample and 36 most specific proteins because it best recovered major cell types and more clearly identified rare cell types, **G**. Gene expression of cell type lineage signatures, **H**. Jaccard similarity coefficient that assessed the clustering stability of CTAPs.

**Supplementary Figure 3. Surface protein specificity and selection for integrative analysis.** Kullback-Leibler divergence measured the specificity of each protein across **A**. all cells, **B**. T cells, and **C**. B/plasma cells. Proteins to the left of the red line were chosen for the CCA integration of each set of cells. Canonical correlations for each of the top 20 canonical variates (CVs) from canonical correlation analysis of **D**. all cells, **E**. T cells and **F**. B/plasma cells, respectively.

**Supplementary Figure 4. Gene and protein features that correlated with the top 20 CVs for integrative analysis.** Correlation z scores for genes (top) and proteins (bottom) in A. T cells, and B. B/plasma cells.

**Supplementary Figure 5. The single-cell CITE-seq RA reference serves as an RA atlas to query other cells. A-B.** We used Symphony^37^ to map synovial cells from the AMP phase I RA dataset (Zhang, *et al.*, 2019) ^21^ onto this AMP phase II single-cell RA reference, **C-D**. We are able to accurately map and predict the cells (Zhang, *et al.*, 2019) from the same cell types with the reference, **E**. We further used Symphony to map cells (Zhang, *et al*, 2019) from each cell type including B/plasma cells (n=1,142), T cells (n=1,529), fibroblasts (n=1,844), and macrophages (n=750) onto the corresponding cell-type specific references from this study (B/plasma cell, T cell, stromal cell, and myeloid cell) to determine correspondence between cell types defined in Zhang, *et al*, 2019 to the cell states in this study. Each heatmap shows results for the major cell type, with rows corresponding to cell states from this study and columns corresponding to cell states from Zhang, *et al*, 2019. Blue-red color scale indicates the log(OR) for a given pair of states (OR is the ratio of odds of mapping a cluster cell in Zhang, *et al*, 2019 to a given cluster of this study compared to odds of mapping other cells in Zhang, *et al*, 2019 onto the same cluster of this study), with higher values indicating greater correspondence between Zhang, *et al.*, 2019 and the fine-grained cell states in this study.

**Supplementary Figure 6. T cell-specific analysis. A.** T cell UMAP colored by fine-grained cell state clusters, **B.** Expression of selected surface proteins among T cells. Cells are colored from blue (low) to yellow (high), **C.** Heatmap of surface protein expression in T cell clusters colored according to the average normalized expression across cells in the cluster, **D.** Heatmap of gene expression in T cell clusters colored according to the average normalized expression across cells in the cluster, scaled for each gene across clusters, **E**. Distribution of T cells across clusters, stratified by CTAP. The size of each segment of each bar corresponds to the average proportion of cells in that cluster across donors from that CTAP. **F.** Number of T cells per donor, stratified by CTAP. Points represent donors. Box plots show median (vertical bar), 25th and 75th percentiles (lower and upper bounds of the box, respectively) and 1.5 x IQR (or minimum/maximum values; end of whiskers).

**Supplementary Figure 7. B/plasma cell-specific analysis. A.** B/plasma cell UMAP colored by fine-grained cell state clusters, **B.** Expression of selected surface proteins among B/plasma cells. Cells are colored from blue (low) to yellow (high), **C.** Heatmap of surface protein expression in B/plasma cell clusters colored according to the average normalized expression across cells in the cluster, **D.** Heatmap of gene expression in B/plasma cell clusters colored according to the average normalized expression across cells in the cluster, scaled for each gene across clusters, **E.** Number of B/plasma cells per donor, stratified by CTAP. Points represent donors. Box plots show median (vertical bar), 25th and 75th percentiles (lower and upper bounds of the box, respectively) and 1.5 x IQR (or minimum/maximum values; end of whiskers), **F.** Heatmap of correlations between select T and B cell subsets, colored by Pearson correlation between per-donor proportions, **G**. Distribution of B/plasma cells across clusters, stratified by CTAP. The size of each segment of each bar corresponds to the average proportion of cells in that cluster across donors from that CTAP.

**Supplementary Figure 8. NK cell-specific analysis. A.** NK cell UMAP colored by fine-grained cell state clusters, **B.** Expression of selected surface proteins among NK cells colored from blue (low) to yellow (high), **C.** Heatmap of surface protein expression in NK cell clusters colored according to the average normalized expression across cells in the cluster, **D.** Heatmap of gene expression in NK cell clusters colored according to the average normalized expression across cells in the cluster, scaled for each gene across clusters, **E**. Distribution of NK cells across clusters, stratified by CTAP. The size of each segment of each bar corresponds to the average proportion of cells in that cluster across donors from that CTAP. **F.** Number of NK cells per donor, stratified by CTAP. Points represent donors. Box plots show median (vertical bar), 25th and 75th percentiles (lower and upper bounds of the box, respectively) and 1.5 x IQR (or minimum/maximum values; end of whiskers).

**Supplementary Figure 9. Myeloid cell-specific analysis. A.** Myeloid cell UMAP colored by fine-grained cell state clusters, **B.** Expression of selected surface proteins among myeloid cells colored from blue (low) to yellow (high), **C.** Heatmap of surface protein expression in myeloid cell clusters colored according to the average normalized expression across cells in the cluster, **D.** Heatmap of gene expression in myeloid cell clusters colored according to the average normalized expression across cells in the cluster, scaled for each gene across clusters, **E**. Distribution of myeloid cells across clusters, stratified by CTAP. The size of each segment of each bar corresponds to the average proportion of cells in that cluster across donors from that CTAP. **F.** Number of myeloid cells per donor, stratified by CTAP. Points represent donors. Box plots show median (vertical bar), 25th and 75th percentiles (lower and upper bounds of the box, respectively) and 1.5 x IQR (or minimum/maximum values; end of whiskers).

**Supplementary Figure 10. Stromal- and endothelial-specific analysis. A.** Stromal cell UMAP colored by fine-grained cell state clusters, **B.** Expression of selected surface proteins among stromal cells colored from blue (low) to yellow (high), **C.** Heatmap of surface protein expression in stromal cell clusters colored according to the average normalized expression across cells in the cluster, **D.** Heatmap of gene expression in stromal cell clusters colored according to the average normalized expression across cells in the cluster, scaled for each gene across clusters, **E**. Distribution of stromal cells across clusters, stratified by CTAP. The size of each segment of each bar corresponds to the average proportion of cells in that cluster across donors from that CTAP, **F.** Number of stromal cells per donor, stratified by CTAP. Points represent donors. Box plots show median (vertical bar), 25th and 75th percentiles (lower and upper bounds of the box, respectively) and 1.5 x IQR (or minimum/maximum values; end of whiskers), **G.** Endothelial cell UMAP colored by fine-grained cell state clusters, **H.** Expression of selected surface proteins among endothelial cells colored from blue (low) to yellow (high), **I.** Heatmap of gene expression in endothelial cell clusters colored according to the average normalized expression across cells in the cluster, scaled for each gene across clusters, **J**. Distribution of endothelial cells across clusters, stratified by CTAP. The size of each segment of each bar corresponds to the average proportion of cells in that cluster across donors from that CTAP. **K.** Number of endothelial cells per donor, stratified by CTAP. Points represent donors. Box plots show median (vertical bar), 25th and 75th percentiles (lower and upper bounds of the box, respectively) and 1.5 x IQR (or minimum/maximum values; end of whiskers).

**Supplementary Figure 11. Clinical and histologic association results using CNA. A**. Representative histologic images illustrating different levels of density and aggregation scores, **B**. For each broad cell type, we identified and presented specific cell populations that were associated with histologic density and aggregation scores by controlling age, sex, and number of cells per sample; cells in red/blue represent positive/negative associations that pass FDR 0.05 correlation, and global permutation p-value is also shown for each association testing.

**Supplementary Figure 12. Association of single-cell RA CTAPs with different clinical characteristics. A**. Clinical, histologic, and ultrasound parameters of patients in each CTAP. For all box plots, each dot represents a donor; boxes show median (vertical bar), 25th and 75th percentiles (lower and upper bounds of the box, respectively) and 1.5 x IQR (or minimum/maximum values; end of whiskers), **B** Association of Krenn inflammation and Krenn lining with CTAPs, adjusting covariates for age, sex, cell number, and clinical collection site. Percent of variance explained by CTAPs only and p-value are calculated with ANOVA test, **C**. CCP levels among seropositive patients alone, **C**. CTAP frequency among seropositive (CCP+, RF+, or both) versus seronegative patients, **D**. CTAP frequency by sex, **E**. CTAP frequency by smoking history, **F**. CTAP frequency by anatomic site of synovial biopsy **H**. Number of patient samples for each CTAP between biopsy and synovectomy, **I.** Collection/cryopreservation sites, **J**. Association of age and RA duration with CTAPs, adjusting covariates for age, sex, cell number, and clinical collection site. Percentage of variance explained by CTAPs alone and p-value are calculated with ANOVA test. 95% confidence intervals are shown. **K**. Sample distributions across CTAPs by recruitment cohort, **L**. Overview of clinical variables for patient samples distributed by CTAPs. “X” represents missing data for a particular sample.

**Supplementary Figure 13. Single-cell cellular sources of cytokines and cytokine receptors.** Z-scored pseudo-bulk expression across the identified 77 cell states of a curated cytokine and receptor list from KEGG (M9809)^105^ is shown. 138 cytokines and receptors that are expressed in more than 3% of total single cells are shown here.

**Supplementary Figure 14. Assigning repeated biopsy and flow samples to CTAPs. A.** Mapping three repeated biopsy samples onto CTAP PC space based on the cell type abundance, **B**. We evaluated CTAP stability by randomly selecting 1,0000 samples and measuring the Mahalanobis distance between these random samples to the baseline samples, **C.** Mapping flow cytometry samples onto CTAP PC space, **D**. Mahalanobis distance of each flow sample to each CTAP centroid; the original CTAP of the single-cell samples from the same donors are labeled as red.

## Supplementary Tables

**Supplementary Table 1. Statistics of demographic, clinical, and histology metrics across recruitment groups and disease activity levels.**

**Supplementary Table 2. Antibodies used in CITE-seq panel.**

**Supplementary Table 3. Antibodies used in flow cytometry panels.**

**Supplementary Table 4**. **Proportions of cell types within each CTAP compared with the proportions across all samples.** For each identified CTAP, we named it based on the cell types if their average proportions were higher in it compared to their average across all samples.

**Supplementary Table 5. Examination of our previously identified RA expanded single-cell clusters**^21^ **in the single-cell dataset from this study.** 95% confidence interval (CI) for the odds ratio (OR) and one-sided MASC (mixed-effects modeling of associations)^106^ p-value are shown.

**Supplementary Table 6. Each identified single-cell cluster’s median proportion across samples within each CTAP.**

**Supplementary Table 7**. **Details and parameters of single-cell integration and clustering for each cell type**. For each broad cell type, we present the number of variable genes, KL divergence threshold for protein selection, Harmony parameters for batch correction, and clustering resolution.

**Supplementary Table 8**. **Differentially expressed genes and relative statistics per CITE-seq cluster.** For each broad cell type, pseudo-bulk differential expression is used with a linear regression model accounting for donor and number of UMIs to identify genes that were more highly expressed inside vs. outside the cluster. Likelihood ratio test p-values and fold change are presented for prioritized markers.

**Supplementary Table 9. Statistics of single-cell cell type-specific associations with CTAPs and histologic parameters.** We show the statistics for each CTAP-specific association testing and histologic parameter association testing.

**Supplementary Table 10. Statistics of demographic, clinical, and histologic metrics across RA CTAPs.** We show the statistics of clinical characteristics, demographic variables, medications, and treatment groups across RA CTAPs.

## Methods

### RA patient recruitment and clinical data collection

The Accelerating Medicines Partnership (AMP) Network for RA and SLE constructed a cross-sectional cohort - samples were collected from 13 clinical sites across the United States and 2 sites in the United Kingdom. The collection occurred over the course of a 45-month period from October 2016 to February of 2020. The study was performed in accordance with protocols approved by the institutional review board. Demographics, RA clinical data, clinical assessments, and measurements of ESR and CRP were performed at the baseline visit. Data collected include age, sex, RA duration, RF or anti-CCP status, RA treatments, tender and swollen joint counts. ESR and CRP were measured using commercial assays in each institution’s clinical laboratory. Disease activity for each subject was calculated using a DAS28-CRP3 validated instrument^91,92^.

### Synovial tissue collection and processing

Synovial tissue samples were obtained from ultrasound-guided biopsies or surgical procedures. Of the 82 samples that completed the tissue processing pipeline, 54 samples were biopsies obtained with a Quick-Core needle, 15 samples were biopsies obtained with portal and forceps, 10 samples were collected during arthroplasty surgery, and 6 samples were collected during surgical synovectomy procedures. All specimens consisted of a median of 13 samples (range 4-36), of which 6-8 fragments were fixed in formalin for subsequent paraffin embedding and processing for histologic analysis. The remaining fragments were cryopreserved in one or more aliquots in Cryostor CS10 (Sigma-Aldrich) cryopreservation media. Samples were shipped to a central biorepository site until sample collection was complete. They were then transited to the central pipeline site, where samples were thawed and processed in batches.

### Histology assessment, definition of density and aggregation for RA synovium

In order to exclude low-quality synovial tissue samples from our multi-omics tissue processing platform, we analyzed hematoxylin and eosin-stained slides of formalin-fixed, paraffin-embedded synovial tissue from each patient. At least six tissue fragments per patient were included in the analysis to mitigate sampling bias. Synovial tissue was identified as previously described^21^, and samples that lacked any discernible synovial tissue were excluded from further analysis. To separate histologic domains of the density of the infiltrate and the extent of formation of discrete aggregates that are not distinguished by the Krenn inflammatory infiltrate domain, we developed consensus semiquantitative four point scales for density and aggregate radial size with a custom atlas using a test set of tissues from the Birmingham Early Arthritis Cohort^93^, scored by three pathologists. This approach was validated by scoring tissues from the first AMP RA cohort^21^, achieving an intra-class correlation coefficient score of 0.896 for summary mean density score of fragments for each tissue and kappa 0.862 for the worst case aggregate score achieved in each tissue. Equivalent ICC figures for the summary mean scores of fragments for Krenn inflammatory domain and Lining layer thickness domains were 0.937 and 0.646 respectively. Three pathologists independently determined Krenn lining and inflammatory infiltrates scores (0-3 each)^94^, cellular density scores (0-3), and aggregate (0-3) scores for each tissue sample, and the mode of the three scores was used for further analysis.

### Tissue disaggregation, live cell sorting, and cell allocations

For pipeline analysis, cryopreserved synovial tissue samples were thawed and disaggregated into single-cell suspension as previously described^95^. Briefly, thawed synovial tissue fragments were mechanically and enzymatically separated in digestion buffer (Liberase TL (Roche) 100 μg/ml and DNase I (New England Biolabs) 100 μg/ml in RPMI) in 37°C water bath for 30 min. Single-cell suspensions from disaggregated synovial tissues were stained with anti-CD235a antibodies (clone 11E4B-7-6 (KC16), Beckman Coulter) and Fixable Viability Dye eFlour 780 (eBioscience/ThermoFisher). Live non-erythrocyte cells (viability dye^-^ CD235^-^) were collected by fluorescence-activated cell sorting (BD FACSAria Fusion). Cells were allocated as follows, in order of priority: (1) 60,000 cells for CITE-seq analysis; (2) 50,000 cells for flow cytometry and bulk RNA-seq analysis; (3) remaining cells re-frozen in aliquots of 70,000 - 100,000 cells in CryoStor CS10 for other analyses (e.g. single-cell ATAC-seq and immune cell repertoire studies). Samples with fewer than 60,000 cells were applied to CITE-seq analysis alone.

### Flow cytometry and bulk RNA-seq

Up to 50,000 sorted live synovial cells were stained with the following antibodies to define cell subsets: CD3 (UCHT1), CD4 (OKT4), CD8 (SK1), CD11c (3.9), CD14 (M5E2), CD19 (HIB19), CD27 (M-T271), CD31 (WM59), CD45 (HI30), CD90 (5E10), CD146 (P1H12), HLA-DR (L234), PD-1 (EH12.2H7). All antibodies were purchased from Biolegend, and staining was performed in the presence of Fc block (eBioscience/ThermoFisher, True-Stain Monocyte Blocker (Biolegend), and Brilliant Stain Buffer (BD Bioscience). We collected flow cytometry data in conjunction with fluorescence-activated cell sorting of up to 1,000 B cells (CD45^+^CD3^-^CD14^-^ CD19^+^), fibroblasts (CD45^-^CD31^-^CD146^-^), macrophages (CD45^+^CD3^-^CD14^+^), and T cells (CD45^+^CD3^+^CD14^-^) on a BD FACSAria Fusion cell sorter.

### Single-cell CITE-seq antibody staining, RNA library preparation, and sequencing

Antibody staining using TotalSeq^TM^-A antibodies was performed as per the recommended protocol (BioLegend). Briefly, we first curated a list of surface proteins based on markers of cell states identified in previous RA studies and TotalSeq^TM^-A antibodies available at the time. To identify optimized concentrations of these antibodies for synovial tissue, we conducted a series of pilot studies where we titrated antibodies and measured their staining quality with mean expression (i.e., intensity) and Kullback-Leibler (K-L) divergence (i.e., specificity). We calculated K-L divergence by comparing the distribution across mRNA-defined clusters of cells expressing the protein highly (>85th percentile) versus the null distribution of all cells. If an antibody had low mean staining and low K-L divergence, we removed it from the panel. If it had high mean staining and low K-L divergence, we titrated it at a lower concentration.

After optimizing the panel and final concentrations (**Supplementary Table 2**), we prepared a cocktail of TotalSeq antibodies and centrifuged for 10 min at 14,000G to remove precipitates. Up to 60,000 sorted live synovial cells were pre-incubated with Human TruStain FcX (BioLegend) in Cell Staining Buffer (BioLegend) for 10 minutes prior to the addition of 100 uL of the antibody cocktail. Single-cell RNA-seq for all synovial samples was performed by the BWH Single Cell Genomics Core. After a 30-minute incubation at 4°C, cells were washed twice in the Cell Staining Buffer and resuspended in 0.4% BSA/PBS. After performing a live cell count using Trypan blue, cells were resuspended at 1,000 cells per microliter and a maximum of 15,000 cells were loaded into a Chromium Next GEM Chip G (10x Genomics). For samples with fewer than 15,000 live cells, all cells were loaded into the chip. cDNA and library generation was done according to the manufacturer’s protocol. mRNA libraries were sequenced to an average of 50,000 reads per cell using Illumina Novaseq S4. CITE-seq antibody-derived tag (ADT) libraries were sequenced to an average of 5,000 reads per cell using Illumina Hi-Seq X Ten.

### Single-cell CITE-seq gene expression and protein expression quantification

We quantified mRNA and antibody-derived tag (ADT) unique molecular identifier (UMI) counts using Cell Ranger v3.1.0. First, raw BCL files were demultiplexed using cellranger mkfastq with default parameters to generate FASTQ files. Then, these FASTQ files were aligned to the GRCh38 human reference genome using Cellranger v3.1.0. Gene and ADT reads were quantified simultaneously using cellranger count.

### Quality control of single-cell CITE-seq data

We show each QC step in **Supplementary Figure 1**. Specifically, we performed consistent QC to remove cells that expressed fewer than 500 genes or contained more than 20% of their total UMIs mapping to mitochondrial genes, resulting in 403,596 cells. Then, we performed sample-level QC and removed samples with a low percentage (< 40%) of cells passing QC. We removed three lower-quality samples (processed on the same day) with less than 40% of cells passing QC compared to 71% for other good quality samples. In the end, we obtained 393,344 cells from 82 samples that passed QC.

We identified and removed doublets based on a combined strategy:

1. To detect doublets/multiplets based on gene count, we utilized the Scrublet^96^ framework implemented in Python on each sample. We input the full raw, unnormalized UMI count data into the Scrublet() function with default parameters. We determined the doublet scores and the threshold for doublet detection by using the scrub_doublets() function with the following parameters: min_counts = 2, min_cells = 3, min_gene_variability_pctl = 85, and n_prin_comps = 30. Based on the distribution of modes of simulated doublet gene expression distributions, we set the threshold at 0.66. Based on this threshold, we identified 4.5% of cells as doublets.
2. Using protein expression, we trained an LDA (Linear Discriminant Analysis)-based classifier on non-doublet cells and then predicted the posterior probability of doublets using cell-type-specific antibodies (CD45, CD3, CD14, CD19, CD20, CD56, CD1C, PDPN, CD146), which improved the precision of doublet detection in a multimodal fashion. We obtained 314,030 cells after doublet detection.

To assess the accuracy of protein measurements in CITE-seq, we selected antibodies for surface markers of each cell-type lineage: T cells (CD45 and CD3D), NK cells (CD45, CD56, CD16, and IL17R), B cells and plasma cells (CD45 and CD19), macrophages (CD45 and CD14), classical dendritic cells (cDCs, CD1C), fibroblast (PDPN), mural cells (PDPN and CD146), and endothelial cells (CD146) (**Supplementary Table 2**). For flow cytometry, we used 13 antibodies (**Supplementary Table 3**). We measured the Pearson correlation between the per-donor proportion of cells in each gate across donors. We removed surface proteins with low expression overall.

### mRNA feature normalization, selection, and scaling

#### Global

For each cell, we normalized the expression of each gene with log(1 + UMIs for gene/total UMIs in cell *10,000). Then, we selected the top 1,000 most highly variable genes in each sample based on a variance stabilizing transformation (VST)^97^, which considers overall variance of the transcript per sample. We excluded cell cycle genes from “Seurat::cc.genes” for downstream analysis. We then pooled the most highly variable genes across all samples for a cell type into a data matrix and performed z-score scaling on each gene to have mean=0 and variance=1 across cells.

#### By cell type

We carried out the same normalization, feature selection, and scaling steps as described for the global analysis, but on only the cells of each given cell type.

### Protein feature normalization, selection, and scaling

#### Global

For each cell, we normalized each protein with centered-log ratio (CLR): {*ln*(*x*_1_/*g*(*x*)),…, *ln*(*x_n_*/*g*(*x*))}, where *x* is a vector of protein counts^98^. For each feature, we then performed z-score scaling on each protein to have mean 0 and variance 1 across cells. To improve discrimination of signal and background in visualizations, we corrected for antibody’ background staining by fitting a Gaussian mixture model (with the normalmixEM function from the mixtools R package; k = 2, lambda = 0.5) to the CLR-normalized expression of each protein in each cell type. Then we calculated the mean of the first (lower) Gaussian in each cell type, identified the lowest mean across cell types, and subtracted this value—representing background—from all cells’ expression of the protein (with a lower bound of 0 for any values that would otherwise become negative).

To select variable proteins, we measured Kullback-Leibler (KL) divergence for each protein by comparing the distribution of cells with normalized expression above the 75th percentile for that protein across broad cell-type clusters, versus the distribution of all cells across broad cell-type clusters. For each feature, we then performed z-score scaling on each protein to have mean=0 and variance=1 across cells. We used a KL-divergence threshold of 0.3.

#### By cell type

We carried out the same normalization as described for global analysis, but only on the cells of each given cell type. For T and B/plasma cells only, we conducted protein feature selection and scaling as described for global analysis. We removed proteins expressed in < 1% of cells and selected variable proteins based on KL divergence (computed as described above except using the 85th percentile to define the distribution of protein-expressing cells). Proteins with KL divergence greater than or equal to 0.025 were considered variable.

### A unimodal dimensionality reduction strategy for single-cell gene expression

For cell-type-specific analysis of myeloid cells, fibroblasts/mural cells, endothelial cells, and natural killer cells, we used a unimodal pipeline to reduce the dimensionality of the data based on mRNA expression. For each cell type, we used truncated principal component analysis (PCA) as implemented in the prcomp_irlba function from the irlba R package^99^ and calculated 20 principal components (PCs) based on the scaled mRNA data. We then corrected sample-driven batch effects with the HarmonyMatrix function from the harmony R package^36^ with parameters as specified in **Supplementary Table 7** and projected the cells into two dimensions with UMAP^100^.

### A multi-modal dimensionality reduction strategy for CITE-seq data

For global analysis of all cell types and cell-type-specific analysis of T and B/plasma cells, we used a multi-modal pipeline to integrate mRNA and surface protein expression from the same cells and project the cells into a low dimensional embedding informed by both modalities^101^. After scaling the protein features so that their total variance was equal to the total variance of the mRNA features, we used canonical correlation analysis (CCA) as implemented in the cc function from the CCA R package to calculate canonical variates (CVs)^102^ based on the scaled mRNA and surface protein data; these are projections of cells onto axes defined by maximally correlated linear combinations of genes and surface proteins that capture the greatest amount of shared variance. For further analysis, we selected the top 20 CVs with highest canonical correlations, as defined in the mRNA space. We then corrected sample-driven batch effects with the HarmonyMatrix function from the harmony R package^36^ with parameters and projected the cells into two dimensions with UMAP^100^.

### Graph-based clustering, differential gene expression, and cell type annotation

We then constructed shared nearest neighbor graphs derived from the top 20 CVs/PCs and applied graph-based Louvain clustering^103^ at various resolution levels (0.2, 0.4, 0.6, 0.8, 1.0). We selected optimized resolution values for each cell type (1.2 for T cells, 0.8 for NK cells, 0.6 for myeloid cells, 0.6 for B cells, 0.6 for stromal cells, 0.3 for endothelial cells) to gain the biological interpretations that made the most sense. We incorporated the number of variable genes chosen per sample and parameters for each cell type’s analytical pipeline in **Supplementary Table 7**. In the end, we identified 24 T cell clusters (94,056 cells), 9 B cell clusters (30,697 cells), 14 NK clusters (8,497 cells), 15 myeloid clusters (76,181 cells), 5 endothelial clusters (25,044 cells), and 10 stromal cell clusters (79,555 cells), for a total of 77 clusters.

For each major cell type, we identified differentially expressed mRNA features and surface proteins by comparing cells from one cluster with all the other cells. We collapsed single-cell mRNA and protein expression profiles into pseudo-bulk count matrices by summing the raw UMI counts for each gene or surface protein across all cells from the same donor and cluster. For mRNA, we tested all mRNA features that were detected in more than 100 cells per donor with non-zero UMI counts. For each feature, we normalized counts in each pseudo-bulk sample into counts per million (CPM). Using linear models, we estimated the effect of each cluster for each feature on pseudo-bulk expression accounting for effects from the donor and the number of UMIs for each pseudo-bulk sample. Next, we used likelihood ratio tests (LRT) between two models: one that has the cluster variable, and another that doesn’t have the cluster variable. Finally, we selected a feature to be a cluster marker if it had a fold change greater than 2 and p-value less than FDR 5%, which is *p* < 0.05/(number of tested genes × number of clusters). We repeated a similar analytical pipeline of normalization and scaling, feature selection, multi-modal dimensionality reduction, clustering, and differential expression analysis for T cells (*p* < 1.5×10^-6^), B cells and plasma cells (*p* < 1.9×10^-6^), NK cells (*p* < 1.6×10^-6^), myeloid cells (*p* < 1.8×10^-8^), stromal cells (*p* < 4.3×10^-7^), and endothelial cells (*p* < 1.2×10^-6^), respectively. Furthermore, we annotated each cell-type cluster based on literature. We present cluster-specific marker genes and relative statistics in **Supplementary Table 8**.

### Building and mapping to global and cell-type-specific references

We used the buildReferenceFromHarmonyObj() function from the Symphony^37^ package to build integrated reference atlases for the global and cell-type specific atlases from the Harmony objects. To find concordance between cell types from our previous study^21^ and this study, we used the Symphony mapQuery() function to map the 5,254 scRNA-seq query cells from Zhang *et al*, 2019 onto the global and respective cell-type reference atlases. We predicted reference cell types and states for the query cells using the knnPredict() function with k=5. For the cell-type specific mapping, we excluded reference dendritic cells or mural cells because they were absent in the query. Note that because the gene expression matrices for the reference (this study) and query^21^ datasets were generated using different versions of Gencode (version 19 vs. version 29, respectively), certain genes were named differently between the two datasets (e.g. *IL8* and *CXCL8* are synonyms for the same gene ENSG00000169429). Because the mapping procedure uses overlapping gene names between reference and query, we “synced” the query gene names to the version 29 names using the shared Ensembl IDs (which do not change between Gencode versions) using the Gencode .gtf files. This converted 9,663 query gene names, and the synced expression matrix was used as input to mapping.

### Identification of CTAPs based on single-cell cell-type abundance

We identified six cell-type abundance phenotypes (CTAPs) based on hierarchical clustering on cell-type abundances for each CITE-seq patient sample. The differences across CTAPs are also reflected in the PCA space. We named each CTAP based on the cell types whose average proportions were higher among samples in the CTAP compared to their average across all samples (**Supplementary Table 4**). To assess the stability of CTAPs, 1) We first bootstrapped the patient samples and clustered the resampled dataset, 2) For every original CTAP subgroup, we found the most similar cluster (based on Jaccard similarity) in each resampled clustering and recorded that value, giving us the maximum Jaccard similarity coefficient for each CTAP. The Jaccard similarity coefficient can be a value between 0 and 1, where 1 indicates complete overlap and 0 indicates no overlap between two sets of the clustering results, 3) We repeated the above two steps 1e4 times and calculated the mean Jaccard similarity coefficient. We performed this process on different possible numbers of patient subgroups ranging from 2 to 10, and evaluated the statistical stability retaining in-group similarity. We selected six clusters as CTAPs because they gave us relatively high stability (mean Jaccard similarity coefficient=0.727) and also high granularity of biologically meaningful interpretations.

### Covarying neighborhood analysis (CNA) to identify cell populations associated with patient CTAP membership

We evaluated whether the global RA CTAPs are associated with changes in the relative abundances of cell states within each of our six major cell types, which would indicate that these CTAP groupings reflect both coarse (relative abundance of major cell types) and fine-scale heterogeneity in synovial tissue composition.

For each major cell type, we used CNA^77^ to associate sample-level attributes to the abundances of cell states within that cell type. CNA defines many small cell neighborhoods in the batch-corrected low-dimensional space and stores that fractional abundance of cells from each sample in each neighborhood in a neighborhood abundance matrix (NAM). By decomposing the NAM with principal component analysis, CNA defines NAM-PCs within each cell type that capture axes of heterogeneity defined by groups of neighborhoods whose abundances vary in a coordinated manner. Here, we use CNA to test for associations between sample-level clinical characteristics and the abundance of covarying neighborhood groups. For associations with histologic metrics such as histology density and aggregate scores, we only used samples that passed histology-level QC grades (Grade A and B). We also use CNA to identify neighborhoods that are associated with one CTAP compared to other CTAPs.

To perform CNA, we used the tl.association() function in the cna Python package with default parameters and top four NAM-PCs as inputs, while controlling for the “age”, “sex”, and “number of cells per sample” as covariates. As CNA utilizes a permutation test, we determined a significant association based on a global permutation *p* < 0.05. For visualization of local associations, which indicate the particular neighborhoods driving a found global association, we used the 5% FDR threshold from CNA to determine which neighborhoods featured a locally significant correlation. In violin plots, we plotted this threshold as dotted lines. In UMAP plots, we colored neighborhoods that pass local significance based on the intensity of their correlation, with red indicating a higher positive correlation, while we colored neighborhoods that did not attain local significance as grey. We used a modified version of CNA, available on Github, which included the following features: 1) scaling the variance per neighborhood within the NAM inversely to the sample size of the source sample for that neighborhood’s anchor cell such that total variance across all neighborhoods anchored on cells from the same sample sums to 1, and 2) the addition of a pseudo-count, a small number that was added to each entry in the NAM. Using CNA, we tested associations of cell neighborhoods that are associated with histology, ultrasound, clinical metrics, and also each CTAP group. The statistics are in **Supplementary Table 9.**

### Modeling histologic, clinical, and demographic characteristics using CTAPs

We used linear mixed modeling to model each histologic parameter and clinical demographic variable using single-cell CTAPs. Only samples that passed histology-level QC (Grade A and B) were included to seek an association between molecular-level categories and histologic metrics. Taking histologic density *Y* as an example, we fitted a mixed-effect model for each CTAP with the number of cells per sample as a cell-level fixed effect, age and sex as demographical level fixed effects, and clinical collection site as a random effect covariate:

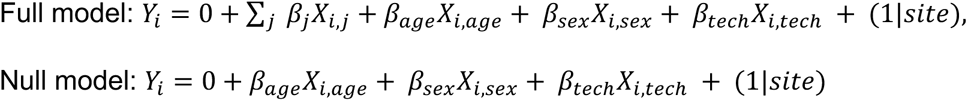

where *β_j_* is the effect size for each CTAP *j* for sample *i*, *β_age_* is a vector of age values and *β_sex_* is a vector of sex values, *β_tech_* is a vector containing a technical covariate that captures the number of cells for each single sample, *X_i_* is the one-hot encoded variable for sample *i* in CTAP *j* as appropriate, and (1|*site*) is the random effect for clinical collection sites. Thus, we used the full model to calculate the corrected values of CTAPs accounting for these technical, cell-level, and donor-level covariates. For modeling age and disease duration, we used a similar model but we removed the age fixed effect from both the full and null model. We obtained percent of variance explained by the CTAPs only by subtracting the variance explained by the null model from the variance explained by the full model. ANOVA p-value was also calculated. The R package lme4 was used for the mixed effect modeling^104^.

### Classifying flow cytometry samples into RA CTAPs

We provided a proof-of-concept framework to assign RA samples processed by other data modalities (e.g., flow cytometry) to the RA CTAPs generated from single-cell technology. Specifically for **Figure 6E**, 1) we quantified the major cell type abundances in a sample using flow cytometry based on cell type markers derived from the single-cell technology, then 2) we mapped each flow sample to the principal component space generated from the CTAP single-cell cell type abundance based on the same features. Here, the features are T, B, Myeloid, stromal, endothelial, and NK cell canonical markers. Now that each flow sample has a loading in the original single-cell abundance space, 3) we built a Mahalanobis-distance-based nearest-neighbor classifier to measure the distance of a flow sample to each of the CTAP centroids. We use Mahalanobis distance to handle the covariance, because our CTAP clusters in PC space are elliptical shaped covariances rather than circular shapes. 4) For each flow sample, we assigned a CTAP label based on which CTAP centroid had the smallest Mahalanobis distance. We calculated the accuracy of our classifications based on a subset (n=15) of RA synovial tissues sent to both single-cell CITE-seq and flow cytometry.

## Acknowledgements

This work was supported by the Accelerating Medicines Partnership (AMP) in Rheumatoid Arthritis and Lupus Network. AMP is a public-private partnership (AbbVie Inc., Arthritis Foundation, Bristol-Myers Squibb Company, Foundation for the National Institutes of Health, GlaxoSmithKline, Janssen Research and Development, LLC, Lupus Foundation of America, Lupus Research Alliance, Merck Sharp & Dohme Corp., National Institute of Allergy and Infectious Diseases, National Institute of Arthritis and Musculoskeletal and Skin Diseases, Pfizer Inc., Rheumatology Research Foundation, Sanofi and Takeda Pharmaceuticals International, Inc.) created to develop new ways of identifying and validating promising biological targets for diagnostics and drug development. Funding was provided through grants from the National Institutes of Health (UH2-AR067676, UH2-AR067677, UH2-AR067679, UH2-AR067681, UH2-AR067685, UH2-AR067688, UH2-AR067689, UH2-AR067690, UH2-AR067691, UH2-AR067694, and UM2-AR067678). Accelerating Medicines Partnership and AMP are registered service marks of the U.S. Department of Health and Human Services. This work was also supported by a Rheumatology Research Foundation Investigator Award and Arthritis National Research Foundation award (to A.H.J.); NIH NHGRI T32HG002295 and NIAMS T32AR007530 (to A. Nathan); NIH NIAMS K08AR077037, Rheumatology Research Foundation Innovative Research Award, and Burroughs Wellcome Fund Career Award in Medical Sciences (to K. Wei); NIH NIGMS T32GM007753 (to J.B.K.); NIH NIAID T32AR007258 (to K.S.); Research into Inflammatory Arthritis Centre Versus Arthritis (22072), IMI-RTCure (777357) and the NIHR Birmingham Biomedical Research Centre (BRC-1215-20009) (to A.F. and D.S.T.); NIH NIAMS K08AR072791 and Burroughs Wellcome Fund Career Award in Medical Sciences (to D.A.R.); NIH NIAID R01AI148435 (to L.T.D.); NIH NIAMS R21AR071670 and P30 AR069655 (to J.H.A.); NIH NIAMS R01AR073833 and R01AR073290 (to M.B.B.); NIH NHGRI U01HG009379 and NIAMS R01AR063759 (to S.R.). We especially acknowledge people in the AMP RA/SLE Network: Arnon Arazi, Celine Berthier, Jill Buyon, Maria Dall’Era, Anne Davidson, Betty Diamond, Andrea Fava, Jennifer Grossman, Nir Hacohen, David Hildeman, Jeffrey Hodgin, Tiffany Hwang, Mariko Ishimori, Ken Kalunian, Diane Kamen, Matthias Kretzler, Holden Maecker, Rong Mao, Maureen McMahon, Fernanda Payan-Schober, Michelle Petri, Chaim Putterman, Daimon Simmons, Thomas Tuschl, David Wofsy, Steve Woodle, and Aaron Wyse.

## Author contributions

L.G-P., K.D.D., D.T., A.C., G.S.F., M.M., I.S., A.B-A., A.M.M., A. Nerviani, F.R., C.P., L.B.H., and D.H., recruited patients and obtained synovial tissues. L.W.M., S.M.G., H.P., V.M.H., A.F., V.P.B., and J.H.A. contributed to the procurement and processing of samples and design of the AMP study. E.D., E.M.G., and B.F.B., performed histological assessment of tissues. D.W., K.P.L., A.F., and V.P.B. curated and analyzed histologic and clinical data. W.A. provided project management and curated histologic and clinical data. K. Wei, A.H.J, G.F.M.W., A. Nathan, and M.B.B. designed and implemented the tissue disaggregation, cell sorting, and single-cell sequencing pipeline. A.H.J., K. Wei, and G.F.M.W supervised and executed the tissue disaggregation pipeline. F.Z., A. Nathan, N.M., Q.X., M.G-A., J.B.K, K. Weinand, J.M., L.R., and S.R. conducted computational and statistical analysis. A.H.J., K. Wei, M.B.B., J.H.A., L.T.D., D.A.R., F.Z., A. Nathan, S.R., D.E.O., J.R-M., and A.F. provided input on cellular analysis and interpretation. D.E.O., J.R-M., A.F., and J.H.A. provided input on histologic analyses. N.M. and K.S. implemented the website. S.R., M.B.B., J.H.A., L.T.D., and D.A.R. supervised the research. F.Z., A.H.J., A. Nathan, N.M., Q.X., and S.R. wrote the initial draft. F.Z., A.H.J., A. Nathan, K. Wei, N.M., D.A.R, L.T.D. J.H.A, M.B.B., and S.R. edited the draft. AMP: RA/SLE Network members contributed to this work by managing patient recruitment, curating clinical data, obtaining and processing synovial tissue samples, managing biorepositories, conducting histological or computational analysis, providing software code, providing website support, and/or providing input on data analysis and interpretation. All authors participated in editing the final manuscript.

## Data availability

All raw and processed data will be available upon acceptance. A cell browser website will be available to visualize our data and results.

## Code availability

All source code will be available on Github upon acceptance. Supplementary Information is available for this paper.

## References

1. Alamanos, Y., Voulgari, P. V. & Drosos, A. A. Incidence and prevalence of rheumatoid arthritis, based on the 1987 American College of Rheumatology criteria: a systematic review. Semin. Arthritis Rheum. 36, 182–188 (2006).

2. Smolen, J. S., et al. Rheumatoid arthritis. Nature Reviews Disease Primers 4, 1–23 (2018).

3. Koduri, G. et al. Interstitial lung disease has a poor prognosis in rheumatoid arthritis: results from an inception cohort. Rheumatology 49, 1483–1489 (2010).

4. McInnes, I. B. & Schett, G. The pathogenesis of rheumatoid arthritis. N. Engl. J. Med. 365, 2205–2219 (2011).

5. Orr, C. et al. Synovial tissue research: a state-of-the-art review. Nat. Rev. Rheumatol. 13, 463–475 (2017).

6. Kerrigan, S. A. & McInnes, I. B. Reflections on ‘older’ drugs: learning new lessons in rheumatology. Nature Reviews Rheumatology vol. 16 179–183 (2020).

7. Nagy, G. & van Vollenhoven, R. F. Sustained biologic-free and drug-free remission in rheumatoid arthritis, where are we now? Arthritis Res. Ther. 17, 181 (2015).

8. Smolen, J. S. & Aletaha, D. Rheumatoid arthritis therapy reappraisal: strategies, opportunities and challenges. Nat. Rev. Rheumatol. 11, 276–289 (2015).

9. Alivernini, S., Laria, A., Gremese, E., Zoli, A. & Ferraccioli, G. ACR70-disease activity score remission achievement from switches between all the available biological agents in rheumatoid arthritis: a systematic review of the literature. Arthritis Res. Ther. 11, R163 (2009).

10. Viatte, S. & Barton, A. Genetics of rheumatoid arthritis susceptibility, severity, and treatment response. Semin. Immunopathol. 39, 395–408 (2017).

11. Amariuta, T., Luo, Y., Knevel, R., Okada, Y. & Raychaudhuri, S. Advances in genetics toward identifying pathogenic cell states of rheumatoid arthritis. Immunol. Rev. 294, 188–204 (2020).

12. Terao, C. et al. Distinct HLA Associations with Rheumatoid Arthritis Subsets Defined by Serological Subphenotype. Am. J. Hum. Genet. 105, 880 (2019).

13. Pitzalis, C., Choy, E. H. S. & Buch, M. H. Transforming clinical trials in rheumatology: towards patient-centric precision medicine. Nat. Rev. Rheumatol. 16, 590–599 (2020).

14. Han, B. et al. Fine mapping seronegative and seropositive rheumatoid arthritis to shared and distinct HLA alleles by adjusting for the effects of heterogeneity. Am. J. Hum. Genet. 94, 522–532 (2014).

15. Oliver, J., Plant, D., Webster, A. P. & Barton, A. Genetic and genomic markers of anti-TNF treatment response in rheumatoid arthritis. Biomark. Med. 9, 499–512 (2015).

16. Smolen, J. S. et al. EULAR recommendations for the management of rheumatoid arthritis with synthetic and biological disease-modifying antirheumatic drugs: 2019 update. Ann. Rheum. Dis. 79, 685–699 (2020).

17. Fraenkel, L. et al. 2021 American College of Rheumatology Guideline for the Treatment of Rheumatoid Arthritis. Arthritis Care Res. 73, 924–939 (2021).

18. Aletaha, D. & Smolen, J. S. Diagnosis and Management of Rheumatoid Arthritis: A Review. JAMA 320, 1360–1372 (2018).

19. Lewis, M. J. et al. Molecular Portraits of Early Rheumatoid Arthritis Identify Clinical and Treatment Response Phenotypes. Cell Rep. 28, 2455–2470.e5 (2019).

20. Humby, F. et al. Rituximab versus tocilizumab in anti-TNF inadequate responder patients with rheumatoid arthritis (R4RA): 16-week outcomes of a stratified, biopsy-driven, multicentre, open-label, phase 4 randomised controlled trial. Lancet 397, 305–317 (2021).

21. Zhang, F. et al. Defining inflammatory cell states in rheumatoid arthritis joint synovial tissues by integrating single-cell transcriptomics and mass cytometry. Nat. Immunol. (2019) doi:10.1038/s41590-019-0378-1.

22. Kuo, D. et al. HBEGF+ macrophages in rheumatoid arthritis induce fibroblast invasiveness. Sci. Transl. Med. 11, (2019).

23. Zhang, F. et al. IFN-γ and TNF-α drive a CXCL10+ CCL2+ macrophage phenotype expanded in severe COVID-19 lungs and inflammatory diseases with tissue inflammation. Genome Med. 13, 64 (2021).

24. Rao, D. A. et al. Pathologically expanded peripheral T helper cell subset drives B cells in rheumatoid arthritis. Nature 542, 110–114 (2017).

25. Alivernini, S. et al. Distinct synovial tissue macrophage subsets regulate inflammation and remission in rheumatoid arthritis. Nat. Med. 26, 1295–1306 (2020).

26. Wang, Y. et al. Rheumatoid arthritis patients display B-cell dysregulation already in the naïve repertoire consistent with defects in B-cell tolerance. Sci. Rep. 9, 1–13 (2019).

27. Wei, K. et al. Notch signalling drives synovial fibroblast identity and arthritis pathology. Nature 582, 259–264 (2020).

28. Bocharnikov, A. V., et al. PD-1hiCXCR5-T peripheral helper cells promote B cell responses in lupus via MAF and IL-21. JCI Insight 4, (2019).

29. Christophersen, A. et al. Distinct phenotype of CD4+ T cells driving celiac disease identified in multiple autoimmune conditions. Nat. Med. 25, 734–737 (2019).

30. Ekman, I. et al. Circulating CXCR5-PD-1hi peripheral T helper cells are associated with progression to type 1 diabetes. Diabetologia 62, 1681–1688 (2019).

31. Martin, J. C. et al. Single-Cell Analysis of Crohn’s Disease Lesions Identifies a Pathogenic Cellular Module Associated with Resistance to Anti-TNF Therapy. Cell 178, 1493–1508.e20 (2019).

32. Na, Y. R., Stakenborg, M., Seok, S. H. & Matteoli, G. Macrophages in intestinal inflammation and resolution: a potential therapeutic target in IBD. Nat. Rev. Gastroenterol. Hepatol. 16, 531–543 (2019).

33. Kochi, Y. Genetics of autoimmune diseases: perspectives from genome-wide association studies. Int. Immunol. 28, 155–161 (2016).

34. Matzaraki, V., Kumar, V., Wijmenga, C. & Zhernakova, A. The MHC locus and genetic susceptibility to autoimmune and infectious diseases. Genome Biol. 18, 76 (2017).

35. Krenn, V. et al. Grading of chronic synovitis--a histopathological grading system for molecular and diagnostic pathology. Pathol. Res. Pract. 198, 317–325 (2002).

36. Korsunsky, I. et al. Fast, sensitive and accurate integration of single-cell data with Harmony. Nat. Methods (2019) doi:10.1038/s41592-019-0619-0.

37. Kang, J. B. et al. Efficient and precise single-cell reference atlas mapping with Symphony. Nat. Commun. 12, 1–21 (2021).

38. Ehrenstein, M. R. et al. Compromised function of regulatory T cells in rheumatoid arthritis and reversal by anti-TNFalpha therapy. J. Exp. Med. 200, 277–285 (2004).

39. Dominguez-Villar, M., Baecher-Allan, C. M. & Hafler, D. A. Identification of T helper type 1-like, Foxp3+ regulatory T cells in human autoimmune disease. Nat. Med. 17, 673–675 (2011).

40. MacDonald, K. G. et al. Regulatory T cells produce profibrotic cytokines in the skin of patients with systemic sclerosis. J. Allergy Clin. Immunol. 135, 946–955.e9 (2015).

41. McClymont, S. A. et al. Plasticity of human regulatory T cells in healthy subjects and patients with type 1 diabetes. J. Immunol. 186, 3918–3926 (2011).

42. Johnson, J. L. et al. The Transcription Factor T-bet Resolves Memory B Cell Subsets with Distinct Tissue Distributions and Antibody Specificities in Mice and Humans. Immunity 52, 842–855.e6 (2020).

43. Wang, S. et al. IL-21 drives expansion and plasma cell differentiation of autoreactive CD11c hi T-bet+ B cells in SLE. Nat. Commun. 9, 1758 (2018).

44. Jenks, S. A. et al. Distinct Effector B Cells Induced by Unregulated Toll-like Receptor 7 Contribute to Pathogenic Responses in Systemic Lupus Erythematosus. Immunity 49, 725–739.e6 (2018).

45. Yuseff, M.-I. et al. Polarized secretion of lysosomes at the B cell synapse couples antigen extraction to processing and presentation. Immunity 35, 361–374 (2011).

46. Rubtsov, A. V. et al. CD11c-Expressing B Cells Are Located at the T Cell/B Cell Border in Spleen and Are Potent APCs. J. Immunol. 195, 71–79 (2015).

47. Radtke, D. & Bannard, O. Expression of the Plasma Cell Transcriptional Regulator Blimp-1 by Dark Zone Germinal Center B Cells During Periods of Proliferation. Front. Immunol. 9, 3106 (2018).

48. Palm, A.-K. E. & Kleinau, S. Marginal zone B cells: From housekeeping function to autoimmunity? J. Autoimmun. 119, 102627 (2021).

49. Weller, S. et al. Human blood IgM ‘memory’ B cells are circulating splenic marginal zone B cells harboring a prediversified immunoglobulin repertoire. Blood 104, 3647–3654 (2004).

50. Tull, T. J. et al. Human marginal zone B cell development from early T2 progenitors. J. Exp. Med. 218, (2021).

51. Descatoire, M. et al. Identification of a human splenic marginal zone B cell precursor with NOTCH2-dependent differentiation properties. J. Exp. Med. 211, 987–1000 (2014).

52. Weyand, C. M. & Goronzy, J. J. Ectopic germinal center formation in rheumatoid synovitis. Ann. N. Y. Acad. Sci. 987, 140–149 (2003).

53. Schröder, A. E., Greiner, A., Seyfert, C. & Berek, C. Differentiation of B cells in the nonlymphoid tissue of the synovial membrane of patients with rheumatoid arthritis. Proc. Natl. Acad. Sci. U. S. A. 93, 221–225 (1996).

54. Dogra, P. et al. Tissue Determinants of Human NK Cell Development, Function, and Residence. Cell 180, 749–763.e13 (2020).

55. Spits, H. et al. Innate lymphoid cells--a proposal for uniform nomenclature. Nat. Rev. Immunol. 13, 145–149 (2013).

56. Ebbo, M., Crinier, A., Vély, F. & Vivier, E. Innate lymphoid cells: major players in inflammatory diseases. Nat. Rev. Immunol. 17, 665–678 (2017).

57. Cella, M. et al. Subsets of ILC3-ILC1-like cells generate a diversity spectrum of innate lymphoid cells in human mucosal tissues. Nat. Immunol. 20, 980–991 (2019).

58. Gordon, S. Phagocytosis: An Immunobiologic Process. Immunity 44, 463–475 (2016).

59. Jakubzick, C. V., Randolph, G. J. & Henson, P. M. Monocyte differentiation and antigen-presenting functions. Nat. Rev. Immunol. 17, 349–362 (2017).

60. Lim, H. Y. et al. Hyaluronan Receptor LYVE-1-Expressing Macrophages Maintain Arterial Tone through Hyaluronan-Mediated Regulation of Smooth Muscle Cell Collagen. Immunity 49, 326–341.e7 (2018).

61. Mulder, K. et al. Cross-tissue single-cell landscape of human monocytes and macrophages in health and disease. Immunity 54, 1883–1900.e5 (2021).

62. Liao, M. et al. Single-cell landscape of bronchoalveolar immune cells in patients with COVID-19. Nat. Med. (2020) doi:10.1038/s41591-020-0901-9.

63. Schuch, K. et al. Osteopontin affects macrophage polarization promoting endocytic but not inflammatory properties. Obesity 24, 1489–1498 (2016).

64. Remmerie, A. et al. Osteopontin Expression Identifies a Subset of Recruited Macrophages Distinct from Kupffer Cells in the Fatty Liver. Immunity 53, 641–657.e14 (2020).

65. Villani, A.-C. et al. Single-cell RNA-seq reveals new types of human blood dendritic cells, monocytes, and progenitors. Science 356, (2017).

66. Zhang, Q. et al. Landscape and Dynamics of Single Immune Cells in Hepatocellular Carcinoma. Cell 179, 829–845.e20 (2019).

67. Korsunsky, I. et al. Cross-tissue, single-cell stromal atlas identifies shared pathological fibroblast phenotypes in four chronic inflammatory diseases. bioRxiv 2021.01.11.426253 (2021) doi:10.1101/2021.01.11.426253.

68. Mizoguchi, F. et al. Functionally distinct disease-associated fibroblast subsets in rheumatoid arthritis. Nat. Commun. 9, 789 (2018).

69. Buechler, M. B. et al. Cross-tissue organization of the fibroblast lineage. Nature 593, 575–579 (2021).

70. Yazici, Y. et al. Efficacy of tocilizumab in patients with moderate to severe active rheumatoid arthritis and a previous inadequate response to disease-modifying antirheumatic drugs: the ROSE study. Ann. Rheum. Dis. 71, 198–205 (2012).

71. Genovese, M. C. et al. Interleukin-6 receptor inhibition with tocilizumab reduces disease activity in rheumatoid arthritis with inadequate response to disease-modifying antirheumatic drugs: the tocilizumab in combination with traditional disease-modifying antirheumatic drug therapy study. Arthritis Rheum. 58, 2968–2980 (2008).

72. Genovese, M. C. et al. Sarilumab Plus Methotrexate in Patients With Active Rheumatoid Arthritis and Inadequate Response to Methotrexate: Results of a Phase III Study. Arthritis Rheumatol 67, 1424–1437 (2015).

73. Wei, K., Nguyen, H. N. & Brenner, M. B. Fibroblast pathology in inflammatory diseases. J. Clin. Invest. 131, (2021).

74. Kalucka, J. et al. Single-Cell Transcriptome Atlas of Murine Endothelial Cells. Cell 180, 764–779.e20 (2020).

75. Wigle, J. T. et al. An essential role for Prox1 in the induction of the lymphatic endothelial cell phenotype. EMBO J. 21, 1505–1513 (2002).

76. Fujimoto, N. et al. Single-cell mapping reveals new markers and functions of lymphatic endothelial cells in lymph nodes. PLoS Biol. 18, e3000704 (2020).

77. Reshef, Y. A. et al. Co-varying neighborhood analysis identifies cell populations associated with phenotypes of interest from single-cell transcriptomics. Nat. Biotechnol. 1–9 (2021).

78. Crotty, S. T Follicular Helper Cell Biology: A Decade of Discovery and Diseases. Immunity 50, 1132–1148 (2019).

79. Dennis, G., Jr et al. Synovial phenotypes in rheumatoid arthritis correlate with response to biologic therapeutics. Arthritis Res. Ther. 16, R90 (2014).

80. Humby, F. et al. Synovial cellular and molecular signatures stratify clinical response to csDMARD therapy and predict radiographic progression in early rheumatoid arthritis patients. Ann. Rheum. Dis. 78, 761–772 (2019).

81. Pitzalis, C., Kelly, S. & Humby, F. New learnings on the pathophysiology of RA from synovial biopsies. Curr. Opin. Rheumatol. 25, 334–344 (2013).

82. Rao, D. A. T Cells That Help B Cells in Chronically Inflamed Tissues. Front. Immunol. 9, 1924 (2018).

83. Seth, A. & Craft, J. Spatial and functional heterogeneity of follicular helper T cells in autoimmunity. Curr. Opin. Immunol. 61, 1–9 (2019).

84. Kroot, E. J. et al. The prognostic value of anti-cyclic citrullinated peptide antibody in patients with recent-onset rheumatoid arthritis. Arthritis Rheum. 43, 1831–1835 (2000).

85. Jansen, L. M. A. et al. The predictive value of anti-cyclic citrullinated peptide antibodies in early arthritis. J. Rheumatol. 30, 1691–1695 (2003).

86. Folkman, J. & D’Amore, P. A. Blood vessel formation: what is its molecular basis? Cell 87, 1153–1155 (1996).

87. Holmes, A. B. et al. Single-cell analysis of germinal-center B cells informs on lymphoma cell of origin and outcome. J. Exp. Med. 217, (2020).

88. Milpied, P. et al. Human germinal center transcriptional programs are de-synchronized in B cell lymphoma. Nat. Immunol. 19, 1013–1024 (2018).

89. Glass, D. R. et al. An Integrated Multi-omic Single-Cell Atlas of Human B Cell Identity. Immunity 53, 217–232.e5 (2020).

90. Nocturne, G. & Mariette, X. B cells in the pathogenesis of primary Sjögren syndrome. Nat. Rev. Rheumatol. 14, 133–145 (2018).

91. Prevoo, M. L. et al. Modified disease activity scores that include twenty-eight-joint counts. Development and validation in a prospective longitudinal study of patients with rheumatoid arthritis. Arthritis Rheum. 38, 44–48 (1995).

92. Wells, G. et al. Validation of the 28-joint Disease Activity Score (DAS28) and European League Against Rheumatism response criteria based on C-reactive protein against disease progression in patients with rheumatoid arthritis, and comparison with the DAS28 based on erythrocyte sedimentation rate. Ann. Rheum. Dis. 68, 954–960 (2009).

93. Choi, I. Y. et al. Stromal cell markers are differentially expressed in the synovial tissue of patients with early arthritis. PLoS One 12, e0182751 (2017).

94. Krenn, V. et al. Synovitis score: discrimination between chronic low-grade and high-grade synovitis. Histopathology 49, 358–364 (2006).

95. Donlin, L. T. et al. Methods for high-dimensonal analysis of cells dissociated from cyropreserved synovial tissue. Arthritis Res. Ther. 20, 139 (2018).

96. Wolock, S. L., Lopez, R. & Klein, A. M. Scrublet: Computational Identification of Cell Doublets in Single-Cell Transcriptomic Data. Cell Syst 8, 281–291.e9 (2019).

97. Stuart, T. et al. Comprehensive Integration of Single-Cell Data. Cell 177, 1888–1902.e21 (2019).

98. Stoeckius, M. et al. Simultaneous epitope and transcriptome measurement in single cells. Nat. Methods 14, 865–868 (2017).

99. Baglama, J. & Reichel, L. Augmented implicitly restarted lanczos bidiagonalization methods. SIAM J. Sci. Comput. 27, 19–42 (2005).

100. McInnes, L., Healy, J. & Melville, J. UMAP: Uniform Manifold Approximation and Projection for Dimension Reduction. arXiv [stat.ML] (2018).

101. Nathan, A. et al. Multimodally profiling memory T cells from a tuberculosis cohort identifies cell state associations with demographics, environment and disease. Nat. Immunol. 22, 781–793 (2021).

102. González, I., Déjean, S., Martin, P. & Baccini, A. CCA: An R Package to Extend Canonical Correlation Analysis. Journal of Statistical Software, Articles 23, 1–14 (2008).

103. Blondel, V. D., Guillaume, J.-L., Lambiotte, R. & Lefebvre, E. Fast unfolding of communities in large networks. arXiv [physics.soc-ph] (2008).

104. Bates, D., Mächler, M., Bolker, B. & Walker, S. Fitting Linear Mixed-Effects Models Using lme4. Journal of Statistical Software, Articles 67, 1–48 (2015).

105. Kanehisa, M., Sato, Y., Kawashima, M., Furumichi, M. & Tanabe, M. KEGG as a reference resource for gene and protein annotation. Nucleic Acids Res. 44, D457–62 (2016).

106. Fonseka, C. Y. et al. Mixed-effects association of single cells identifies an expanded effector CD4+ T cell subset in rheumatoid arthritis. Sci. Transl. Med. 10, (2018).

