## Supplementary figures for "Cellular deconstruction of inflamed synovium defines diverse inflammatory phenotypes in rheumatoid arthritis"

### Supplementary Figure 1

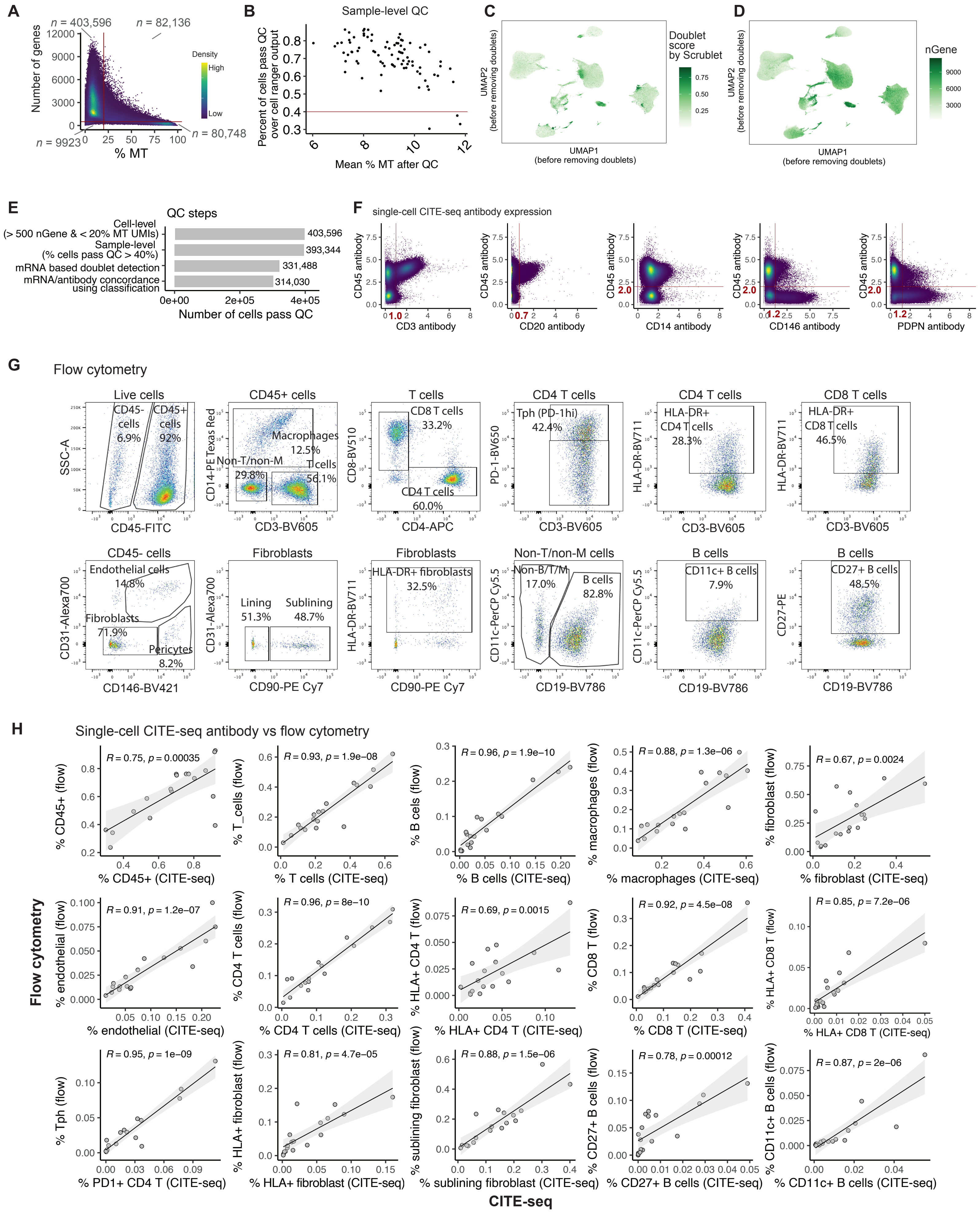

### Supplementary Figure 2

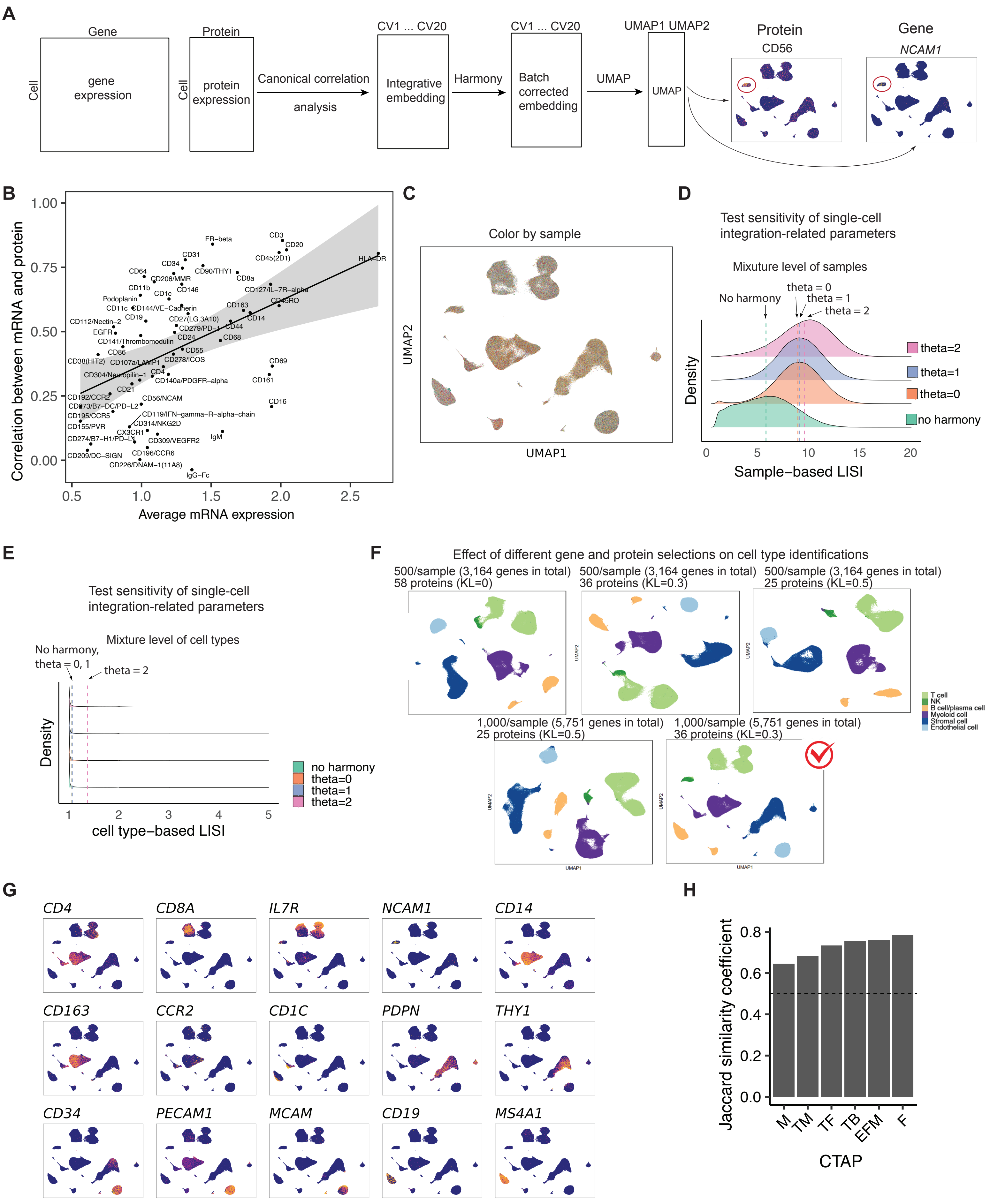

### Supplementary Figure 3

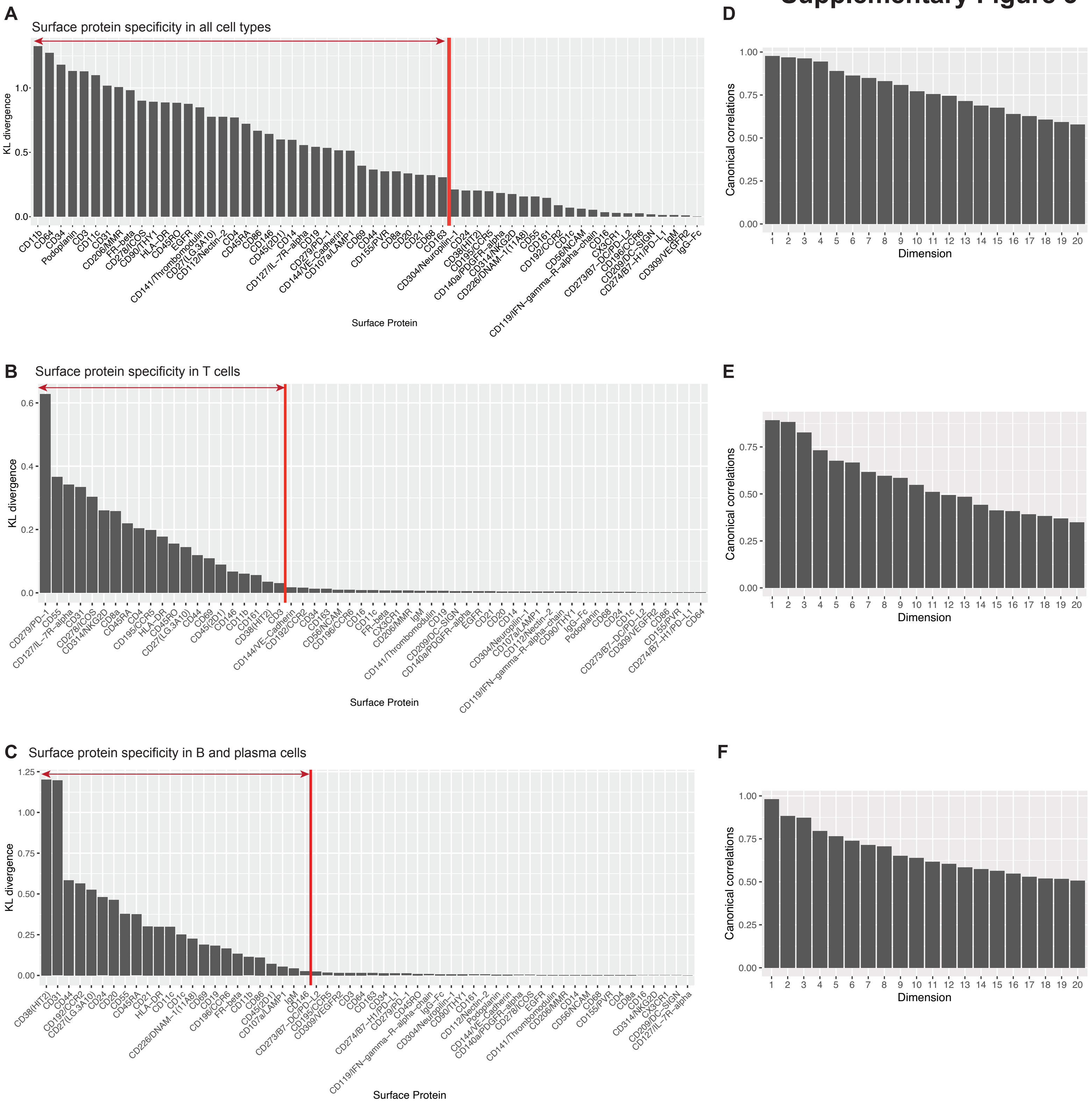

#### Supplementary Figure 4

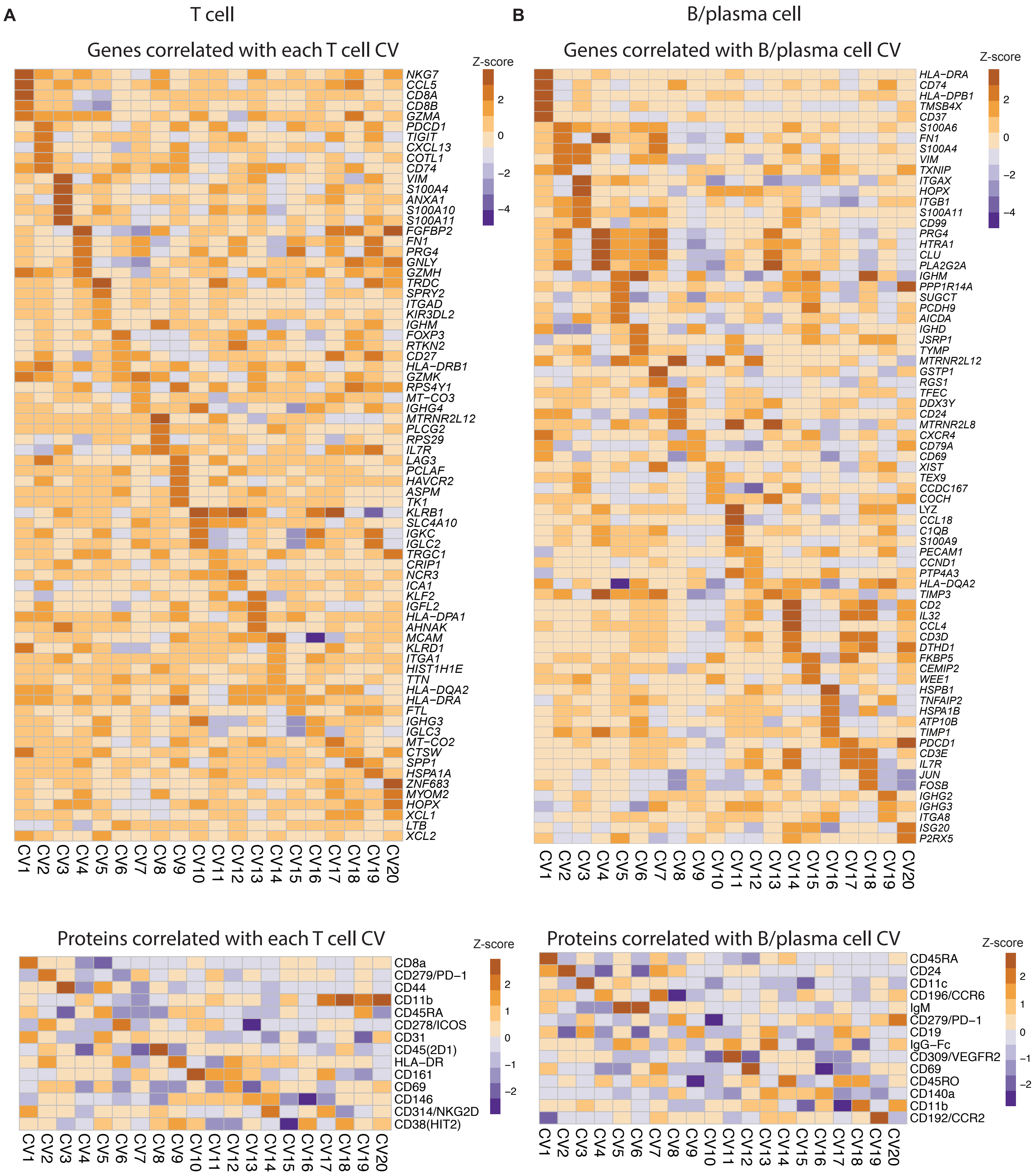

### Supplementary Figure 5

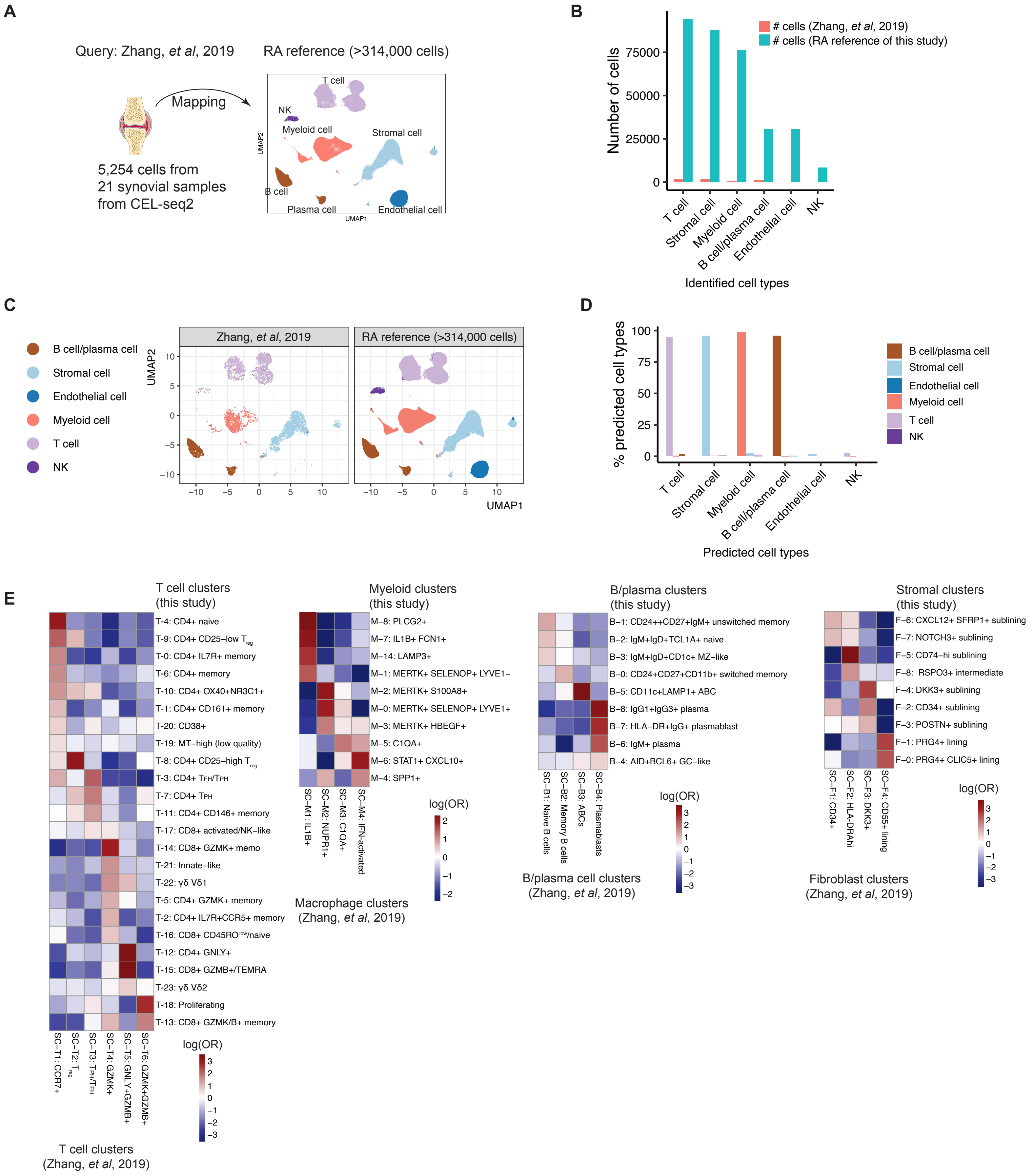

### Supplementary Figure 6

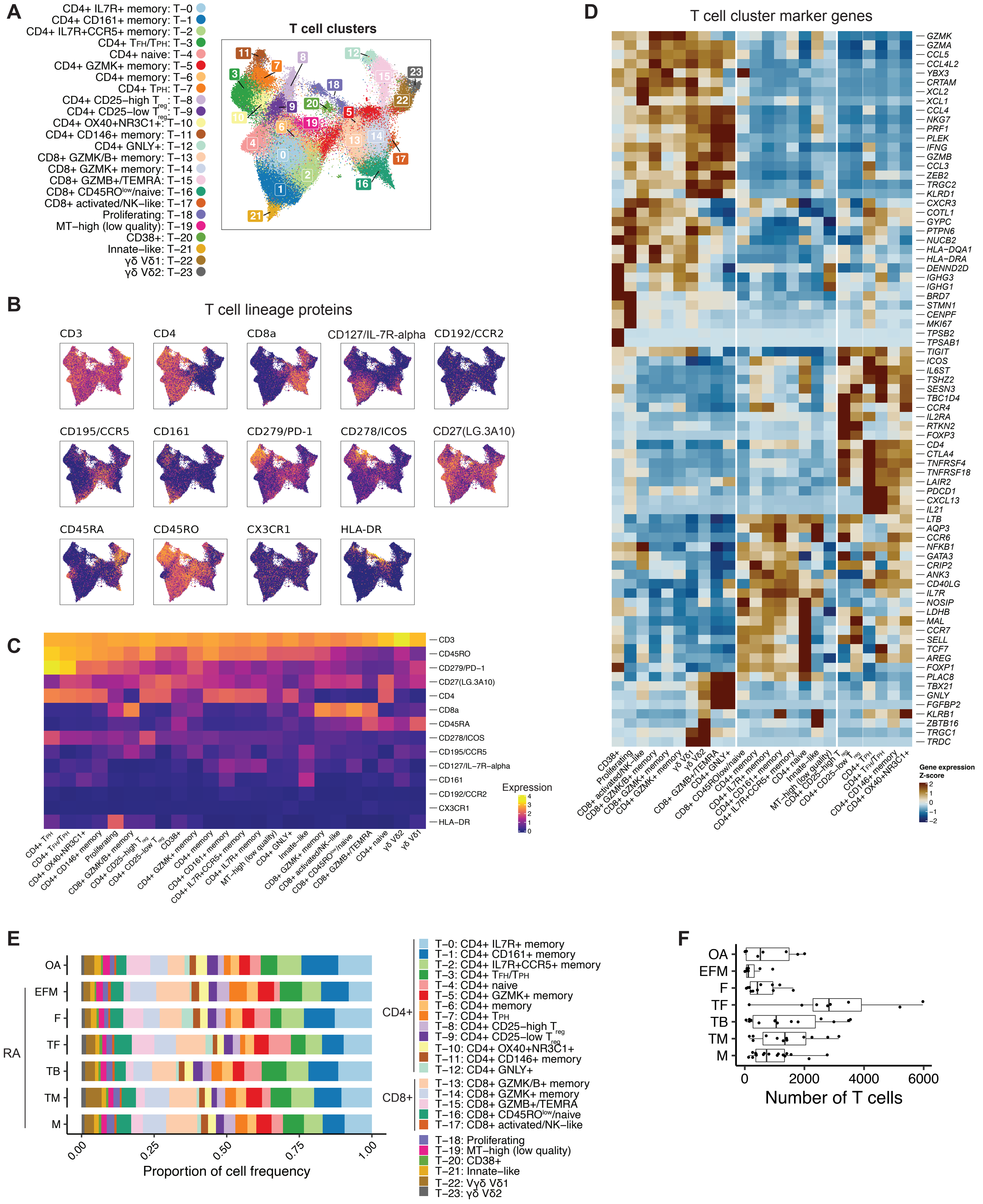

#### Supplementary Figure 7

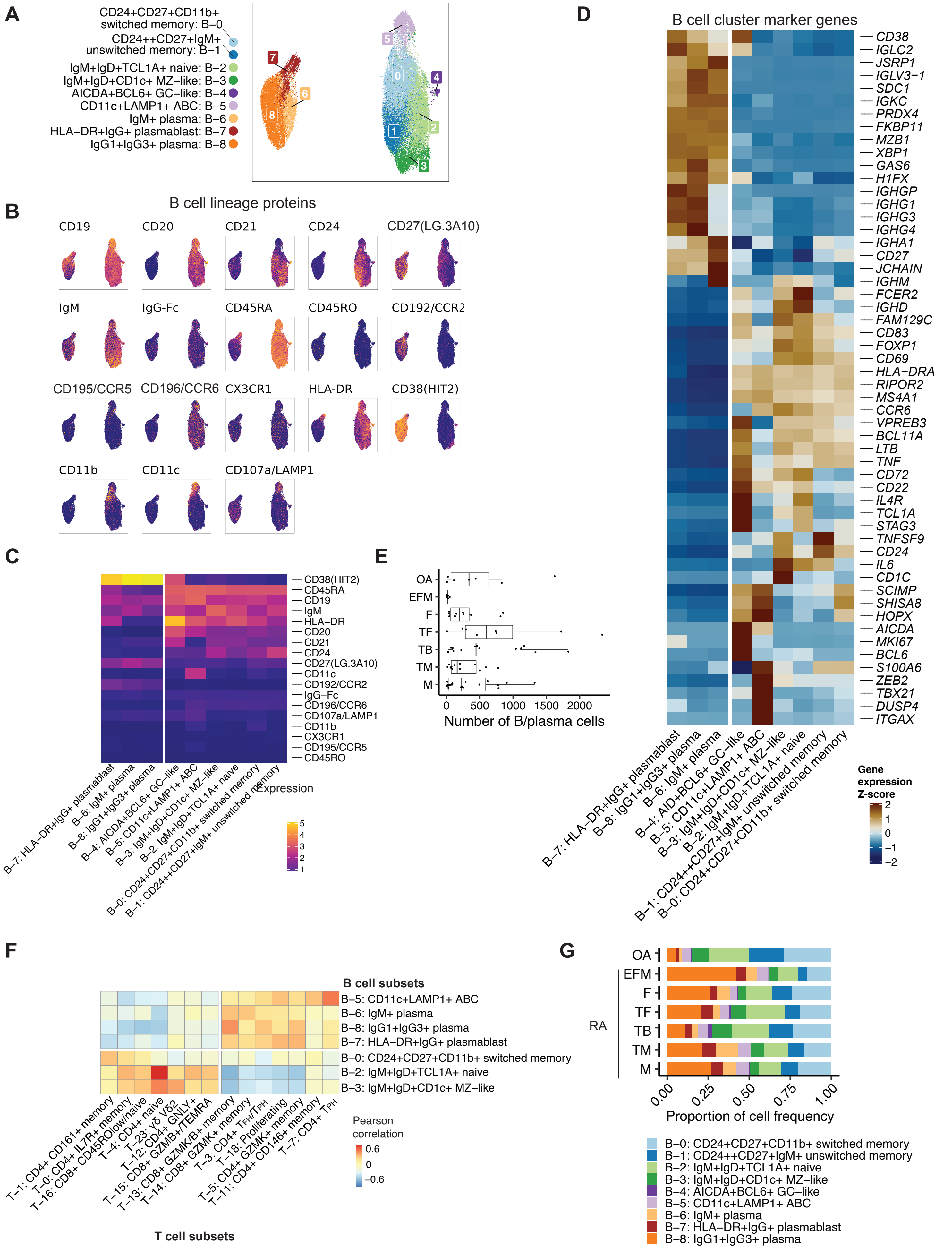

### Supplementary Figure 8

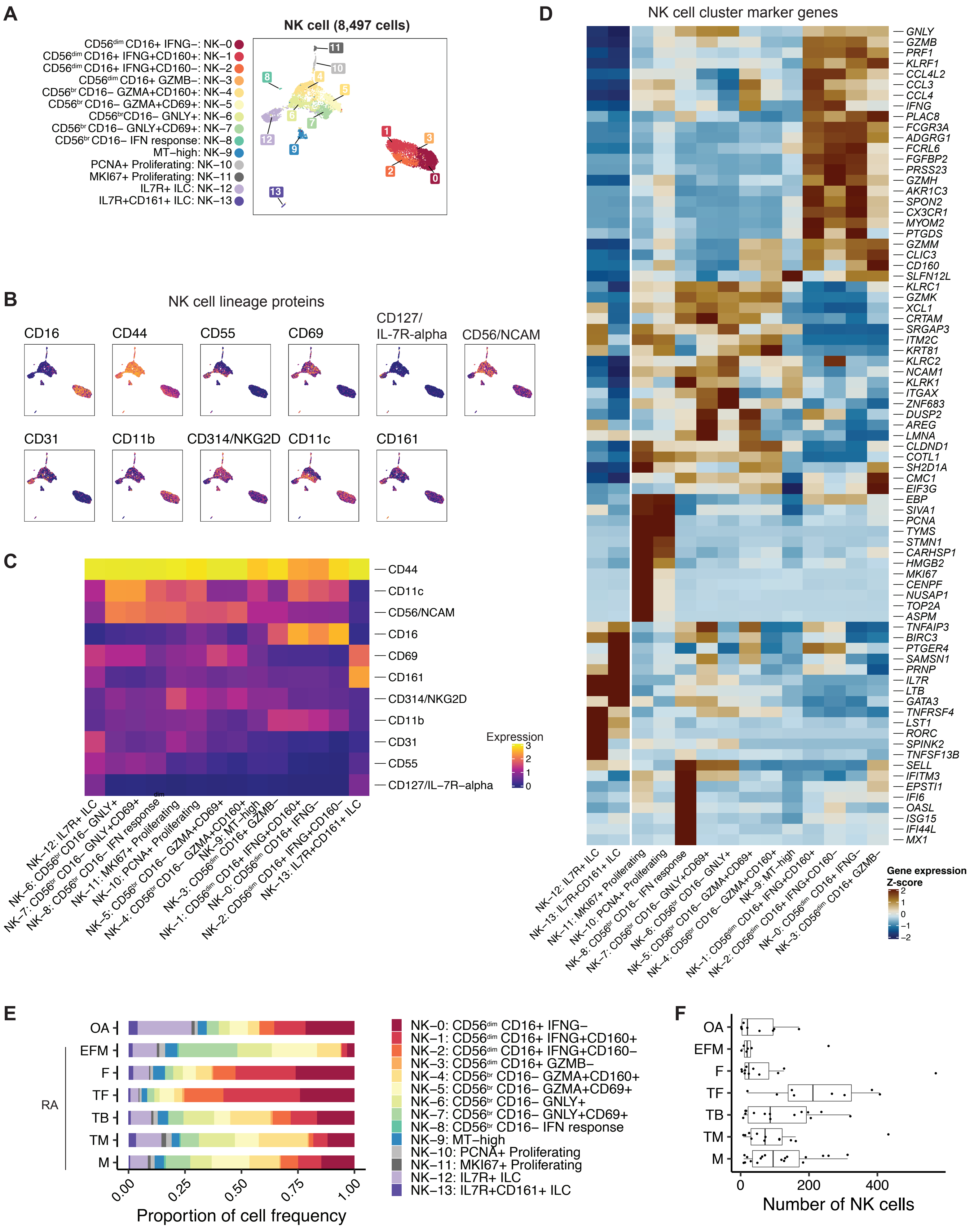

### Supplementary Figure 9

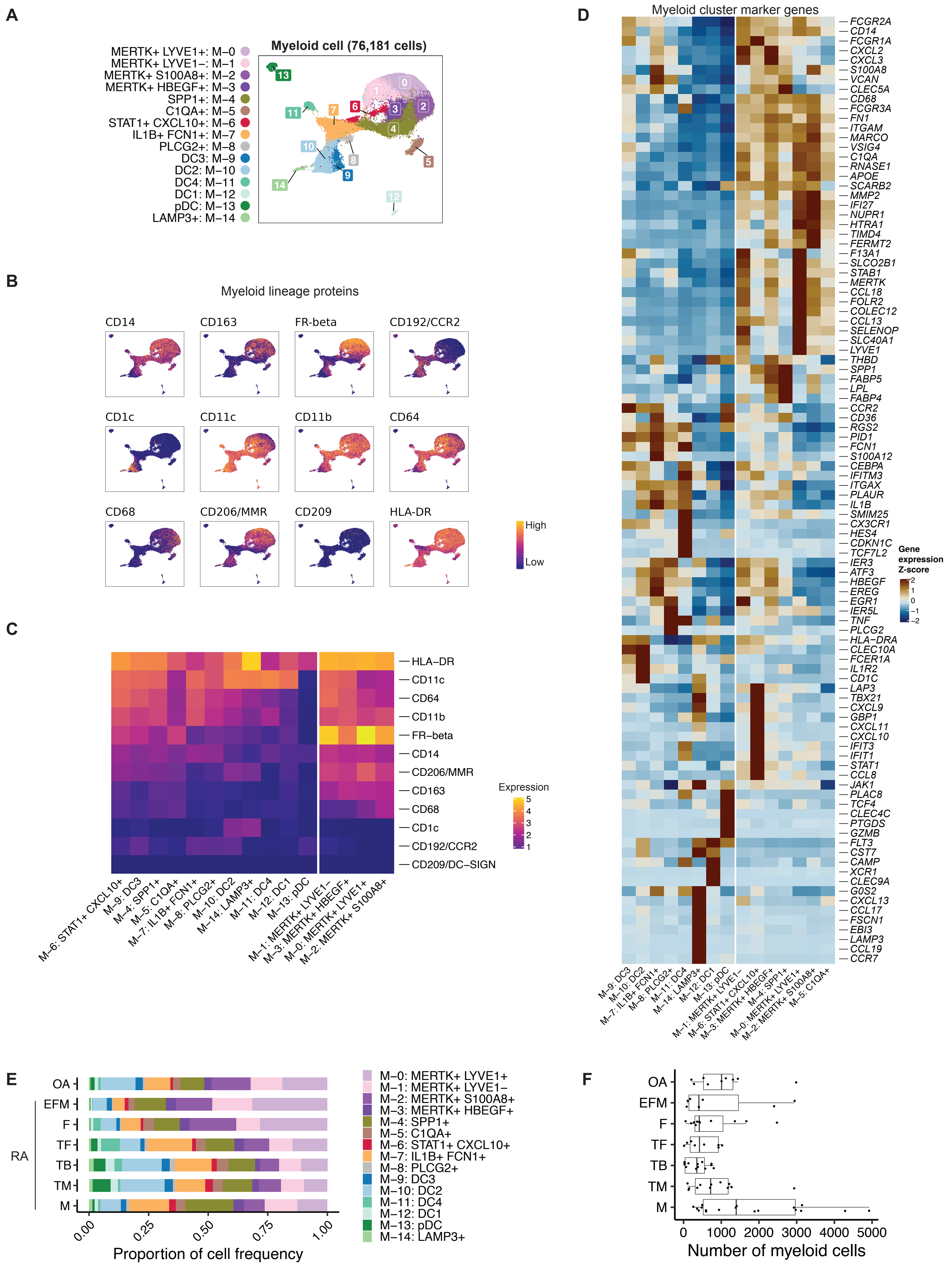

### Supplementary Figure 10

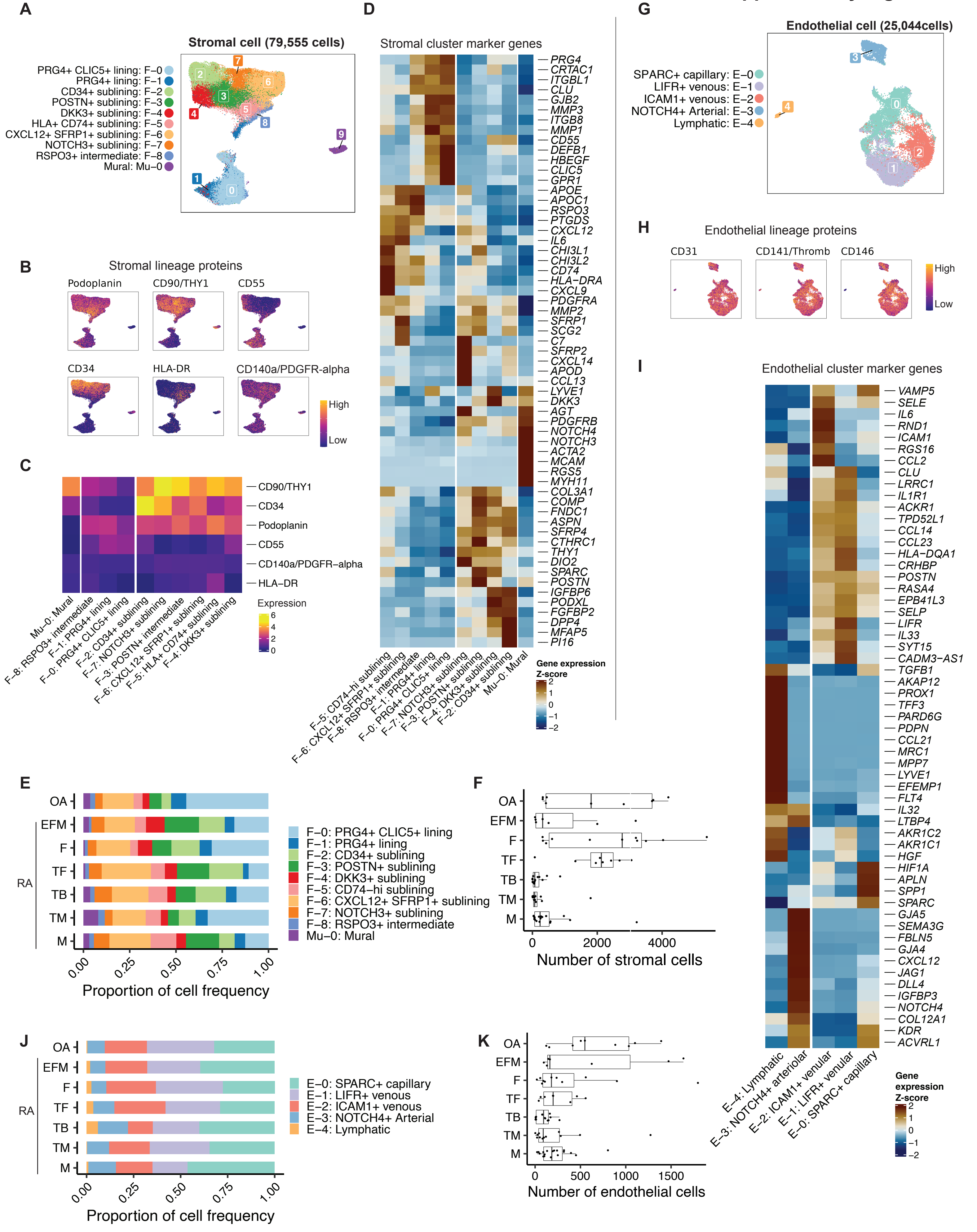

### Supplementary Figure 11

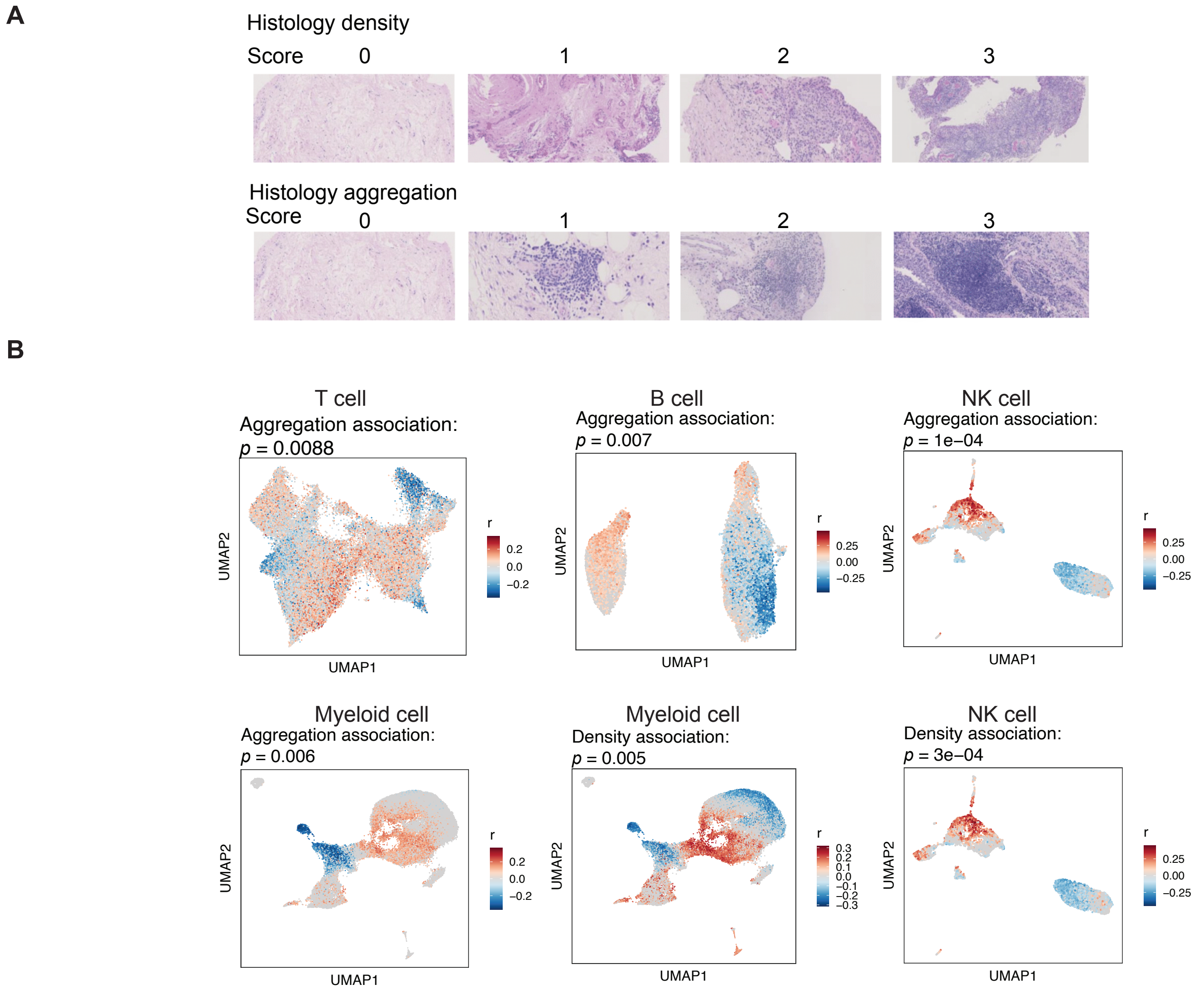

### Supplementary Figure 12

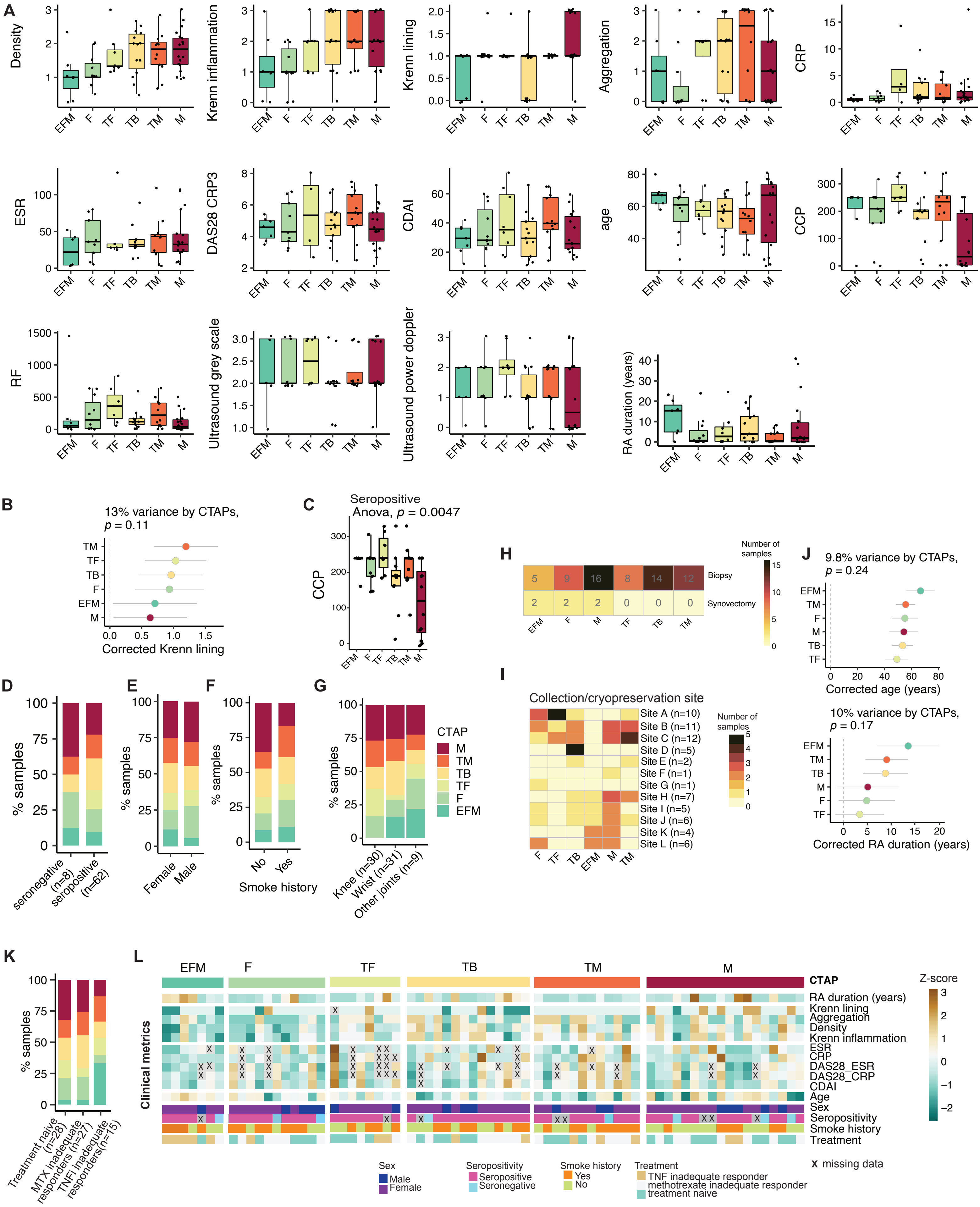

### Supplementary Figure 13

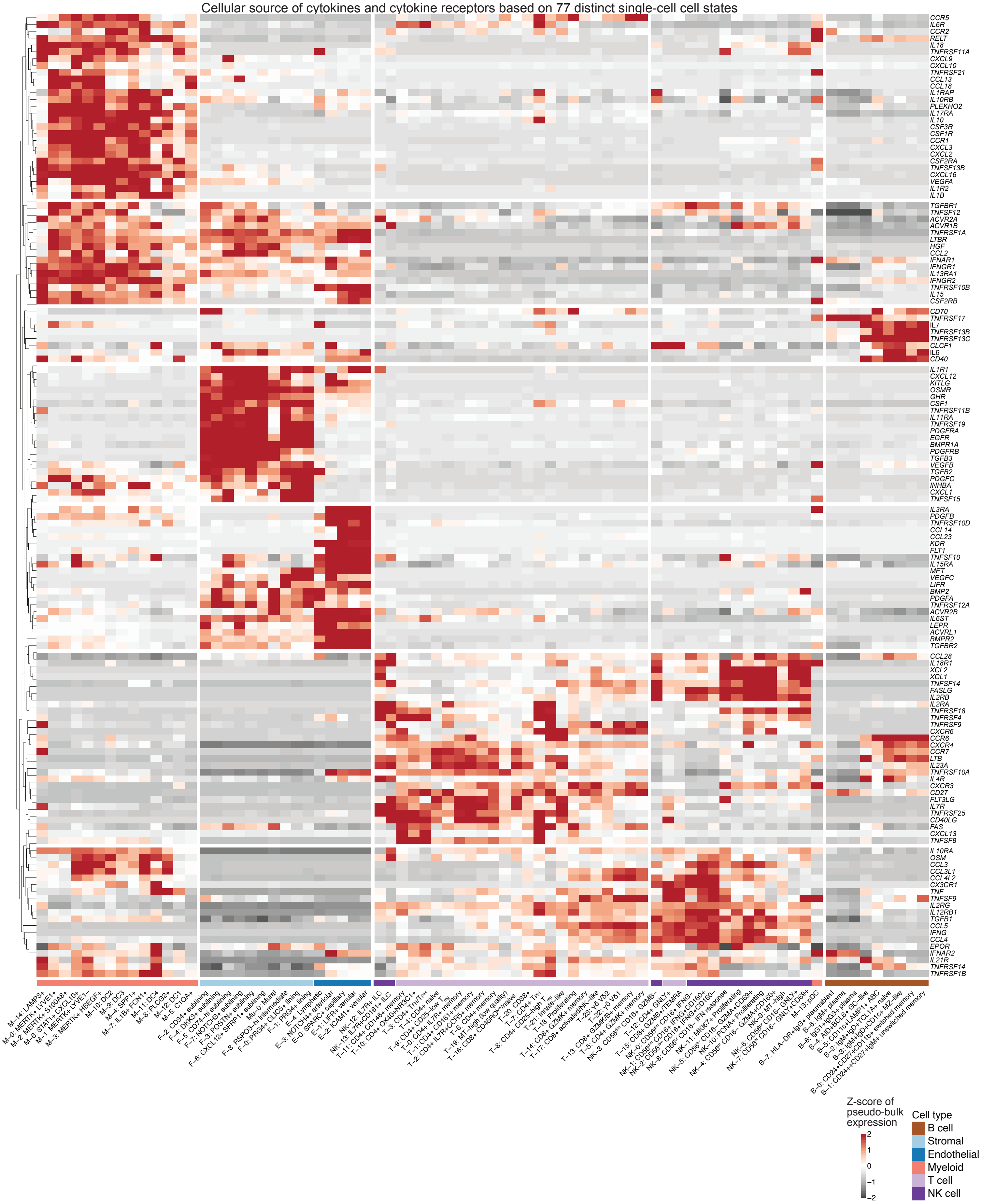

#### Supplementary Figure 14

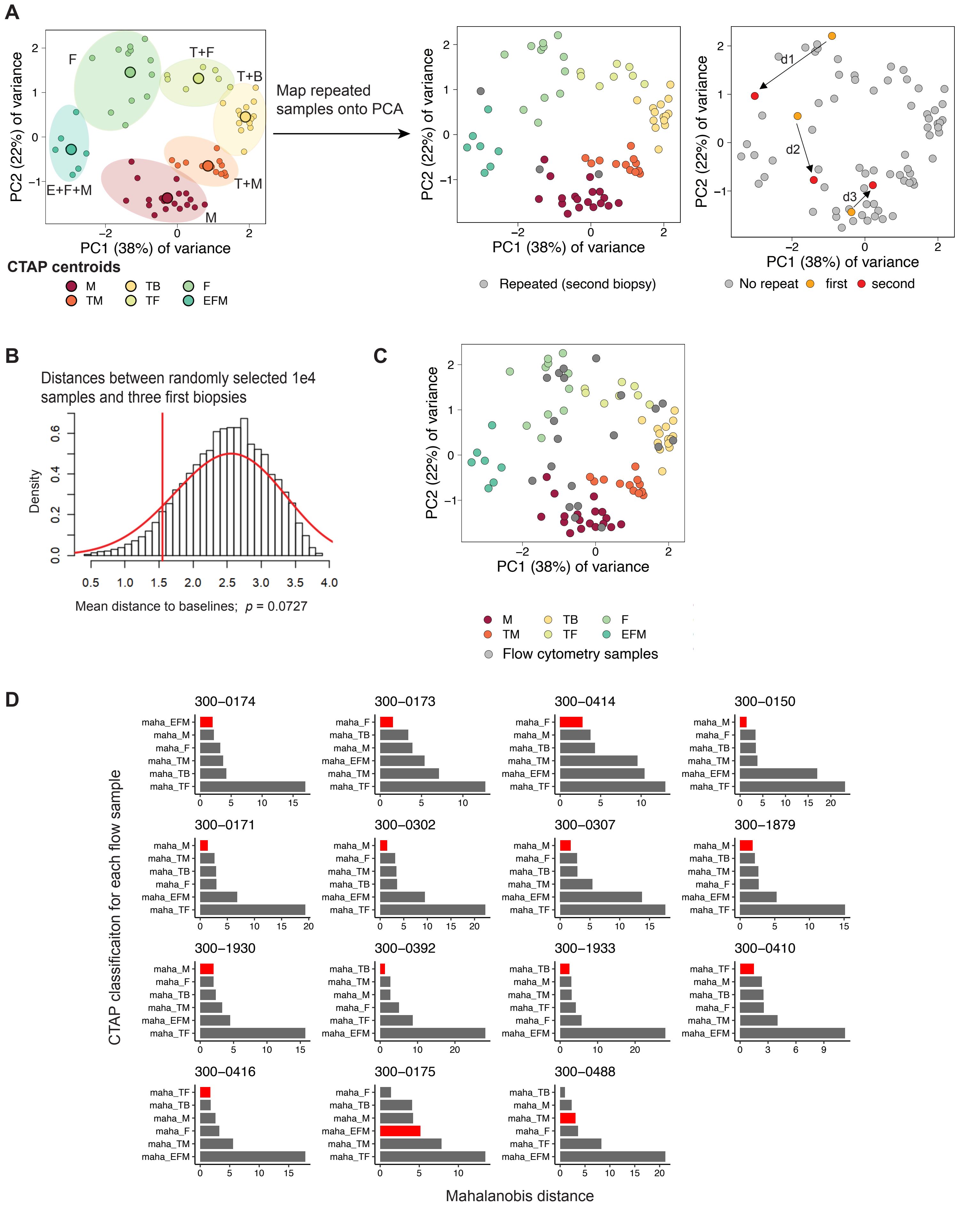
